# Fis family members synergistically control the virulence of *Legionella pneumophila*

**DOI:** 10.1101/2024.09.25.615009

**Authors:** Claire Andréa, Julie Bresson, Christophe Ginévra, Anne Vianney, Nathalie Bailo, Annelise Chapalain, Laetitia Attaiech, Kevin Picq, Caroline Ranquet, William Nasser, Patricia Doublet, Elisabeth Kay

**Affiliations:** CIRI, Centre International de Recherche en Infectiologie, (Team : Legionella pathogenesis), Univ Lyon, Inserm, U1111, Université Claude Bernard Lyon 1, CNRS, UMR5308, ENS de Lyon, F-69007, Lyon, France; CNRL, Centre national de référence des légionelles, centre de biologie et de pathologie nord, institut des agents infectieux, hôpital de la Croix Rousse. 103, grande rue de la Croix Rousse, 69317 Lyon cedex 04, France; CIRI, Centre International de Recherche en Infectiologie, (Team : HORIGENE), Univ Lyon, Inserm, U1111, Université Claude Bernard Lyon 1, CNRS, UMR5308, ENS de Lyon, F-69007, Lyon, France; BGene-Genetics, 7 rue des Arts et Métiers, 38000 Grenoble, France; Laboratoire Microbiologie, Adaptation et Pathogénie, Univ Lyon, CNRS, Unité Mixte de Recherche 5240, INSA Lyon, Université Claude Bernard Lyon 1, 11 avenue Jean Capelle, 69621 Villeurbanne, France

**Keywords:** *Legionella pneumophila*, Nucleoid-associated proteins (NAPs), Factor for Inversion Stimulation (FIS), effectors, Dot/Icm T4SS

## Abstract

*Legionella pneumophila* virulence is controlled in a growth phase-dependent manner by a complex regulatory network involving several two-component systems, small regulatory RNAs and the translational CsrA regulator. Here, we address the additional role of Nucleoid-associated proteins (NAP) regulators in this network, by investigating the regulatory functions of the three Fis paralogs (Fis1, Fis2, Fis3), a unique feature among bacteria, in the infection cycle of *L. pneumophila*. Specifically, we show that deletion of *fis1* has a major impact on *L. pneumophila* virulence, and that deletion of *fis2* enhances the intensity of this phenotype. Consistently, RNA-seq analysis and reporter gene fusions demonstrate the predominant role of Fis1 in the regulation of many virulence-related genes, including those involved in the flagellum, pili biosynthesis, and Dot/Icm type 4 secretion machinery, as well as several genes encoding Dot/Icm effectors. Both Fis1 and Fis2 bind to AT-rich motifs upstream their target genes, but Fis1 with higher affinity than Fis2. Importantly, Fis1 and Fis2 would be capable of forming heterodimers that could bind with variable affinity to this AT-rich motif. It is also important to note that the three Fis proteins are not produced at the same time and in the same amounts. We therefore hypothesize that the duplication of *fis* genes in *L. pneumophila* is not simply a back-up system to compensate for potentially deleterious mutations in a *fis* gene, but rather a means to fine-tune the expression of targeted genes, particularly virulence genes.

**IMPORTANCE:** Appropriate control of virulence gene expression is crucial to the success of bacterial infection. Nucleoid-associated protein regulators, including Fis proteins, have been shown to participate in the virulence of several human pathogens. The importance of our discovery lies in the fact that *L. pneumophila* possesses three non-homologous Fis proteins instead of just one. We demonstrate that Fis1 and Fis2 are not functional duplicates of each other. On the contrary, Fis1 and Fis2 are synthesized neither simultaneously nor in equal amounts during the bacterial growth phase, and they cooperate to regulate virulence gene expression by targeting similar AT-rich motifs, albeit with distinct affinity, and by being capable of forming heterodimers. Taken together, our data suggest that the high conservation of *fis* gene duplication results from the need for fine-tuned control of *Legionella* virulence in response to its different environmental and human hosts, rather than from functional redundancy to circumvent deleterious *fis* mutations.

## INTRODUCTION

Nucleoid-associated proteins (NAPs), such as Fis proteins, are conserved small basic proteins capable of binding and bending DNA, therefore influencing DNA topology and the expression of many genes, either positively or negatively, depending on the location of their binding sites relative to the gene promoter sequence (1, 2). Originally identified as <u>F</u>actor for the Inversion <u>S</u>timulation promoting site-specific recombination reactions catalysed by Hin and Gin DNA recombinases of *Salmonella* and phage Mu respectively (3, 4), Fis has also been shown to play a versatile role in the regulation of many diverse physiological processes, such as the initiation of DNA replication (5), DNA supercoiling (6, 7), the site-specific recombination of bacteriophage lambda (8, 9) and the transcription of rRNA and tRNA operons (10–12). It is also considered as a global transcriptional regulator that participates in the control of quorum sensing (13), biofilm formation (14, 15), antibiotic resistance (16) and virulence of diverse pathogens (17–25). Strikingly, Fis protein is encoded by a single chromosomal copy of the *fis* gene in all bacterial species, with the exception of all *Legionella* species, which have three non-homologous copies of the *fis* gene in their genome. This unique feature among bacteria is presumably the result of two duplication events that occurred prior to the divergence of the *Legionella* genus (26).

*L. pneumophila* is a facultative intracellular parasite able to replicate within many different free-living freshwater *protozoa* and to survive into the extracellular environment until the infection of a new host (27–29). It is also an accidental human pathogen replicating within alveolar macrophages, thus causing a severe pneumonia termed Legionnaires’ disease (30–34).

*L. pneumophila* infects its phagocytic, environmental and human hosts in a similar biphasic cycle, comprising a replicative phase and a transmissive phase. Crucial for establishing this cycle is the Dot/Icm type 4 secretion system (T4SS) (35, 36) which translocates more than 300 effector proteins that modulate host cell functions into infected cells (34, 37–41). Specifically, following phagocytosis, the bacteria escape endocytic degradation and promote the biogenesis of an endoplasmic-reticulum (ER)-derived vacuole, called *Legionella*-containing vacuole (LCV), in which the bacteria replicate (42–46). During intracellular replicative phase, genes involved in bacterial replication and central metabolism are expressed while those involved in virulence, cytotoxicity, stress resistance and motility (also called transmissive traits) are repressed, mainly due to the post-transcriptional control exerted by the RNA-binding protein CsrA (47–49). After extensive multiplication within the LCV, *L. pneumophila* undergoes a global change in gene expression leading to the derepression of transmissive traits in order to promote host cell lysis and the infection of new phagocytic cells (50–52). The switch between replicative and transmissive phases results from nutritional stress, in particular depletion in amino acids and perturbation in fatty acids biosynthesis that leads to the accumulation of the alarmone (p)ppGpp. This accumulation triggers the two-component system (TCS) LetA/LetS (GacA/S orthologues) and increases the amount of the alternative sigma factor of stress response, RpoS. Both transcriptional regulators, LetA and RpoS, are required for the full expression of the regulatory RNAs RsmX, RsmY and RsmZ that bind and sequester the repressor CsrA, thus relieving the expression of transmissive traits and about 40 Dot/Icm effector-encoding genes (47, 53–56). Additional regulatory systems control virulence in *L. pneumophila*, including the *Legionella* Quorum Sensing system LqsST/R (57, 58) and the TCSs PmrA/B (59–61) and CpxR/A (62–66) All these systems were shown to control the expression of both Dot/Icm translocated effectors encoding genes and several *dot/icm* secretion machinery genes, at least for CpxR/A (63, 64) and PmrA /B TCSs (61). PmrA response regulator is also required for CsrA transcription while LqsR expression depends on CsrA activity (47), thus connecting all these regulatory pathways the ones with the others.

Here, we address the question of the role of the three Fis1, Fis2, and Fis3 proteins in the control of the virulence of *L. pneumophila* Paris strain, with a particular attention paid to potential specific, redundant or complementary roles of each copy of *fis* genes. In a previous study, Fis1 and Fis3, but not Fis2, were shown to directly repress the expression of 18 Dot/Icm effector-encoding genes in the Philadelphia strain (26). More recently, three additional effector-encoding genes and their cognate regulators, located on two distinct genomic islands found in only a few *Legionella* species, have also been reported to be silenced by Fis1 and Fis3 (67, 68). In the present study, we demonstrate the predominant role of Fis1 in regulating genes at the onset of the post-exponential/transmissive phase, by directly targeting genes that encode transmissive traits such as flagella and type IV pili, but also genes that encode major players of the complex regulatory network described above. We also show that Fis2 complete this role by synergically targeting the same set of genes, thus demonstrating a novel role of Fis2 in regulating of key virulence factors. We also demonstrate that Fis1 and Fis2 control the genes encoding the components of the Dot/Icm T4SS, as well as many effector genes, thus contributing to the fine regulation of *L. pneumophila*’s main virulence factors. Finally, our data reveal that Fis1 and Fis2, which are synthesized neither simultaneously, nor in equal amounts during bacterial growth phase, bind to the same regulatory sequences on promoter regions of target genes, albeit with distinct affinity. This suggests that Fis2 plays a crucial role in finely regulating the expression of targeted genes, particularly those involved in *L. pneumophila* virulence, rather than merely serving as a redundant copy of Fis1. It’s worth noting that despite our inability to obtain a *fis3* mutant in the Paris strain, probably due to its essentiality in this genetic context, we have obtained insights showing that Fis3 exhibits more polymorphism within the *L. pneumophila* species and the genus *Legionella* and is distinguished by a slightly different structural organization from Fis1 and Fis2, which might suggest a different functional role from its paralogs Fis1 and Fis2.

## RESULTS

### Occurrence and diversity of the three Fis paralogs among *Legionella* species

Zusman et al. (26) demonstrated that the three *Fis* paralogs are present in four distinct *Legionella* species and hypothesized that these paralogs originated from two gene duplication events prior to the divergence of the *Legionella* genus. To further investigate the evolutionary conservation of *Fis* paralogs across the genus, we conducted a comparative genomic analysis examining their presence in 64 non-*Legionella pneumophila* species and 6 *L. pneumophila* strains **(Fig. 1A and Table S1)**. Phylogenetic reconstructions confirm that Fis1, Fis2, and Fis3 are conserved across all species analyzed, with Fis2 and Fis3 displaying greater genetic diversity compared to Fis1. Notably, Fis1 shows a high degree of conservation with an average amino acid identity exceeding 92%, whereas Fis2 and Fis3 exhibit substantial sequence variation, with identities ranging from 73–100% for Fis2 (average 84%) and 72–100% for Fis3 (average 86%). This increased sequence variability of Fis2 and Fis3 suggests that that these paralogs may be under more relaxed selective pressures than Fis1, potentially allowing for adaptive divergence in certain species.

**Fig. 1.**
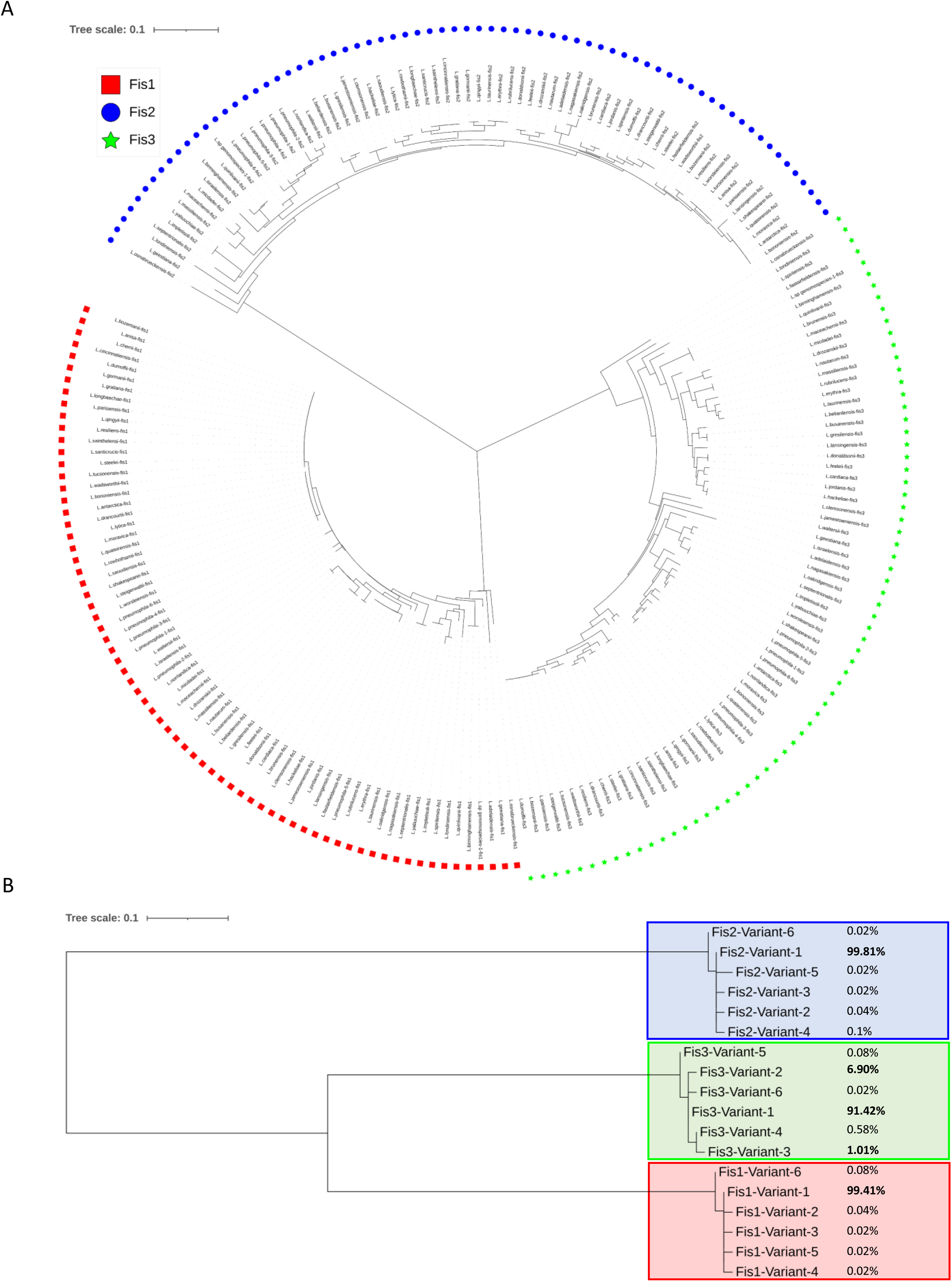
Fis paralog conservation and diversity in *Legionella* species. (A) BioNJ Tree of Fis1, Fis2 and Fis3 from one representative genome of each *Legionella* non-*pneumophila* species (n=64) and one representative sequence of each of *L. pneumophila* fis variants (n=6). Phylogenetic tree represents the phylogenetic diversity of Fis1 (red squares), Fis2 (blue circles) and Fis3 (green stars) (B) PhyML tree of one representative variant of fis1, 2 and 3 extracted from 5216 *Legionella pneumophila* genomes from NCBI. Values on the right of the tree’s leafs correspond to the ratio of each variant in the genome dataset.

To further explore the genetic diversity of Fis paralogs at the species level, we analyzed the protein sequence variability in 5,216 *L. pneumophila* genomes currently available from the NCBI **(Fig. S1)**. Several distinct allelic variants have been identified for each Fis proteins, but the proportion of these variants vary according to the protein **(Fig.1B)**. Fis1 and Fis2 are highly conserved within the *L. pneumophila* species, with at least 99.4% of strains possessing identical protein sequences. The other alleles are present in a very small number of strains (between 1 and 5). In contrast, Fis3 displays higher polymorphism. The predominant allelic variant was found in 91.4% of the strains (4,768 strains), while alleles 2 and 3 were detected in 360 and 52 other strains, respectively. This pattern suggests increased sequence diversity of Fis3 within the *L. pneumophila* population, which could potentially lead to functional variability.

### Deleting *fis1* and *fis2* impairs LCV biogenesis and intracellular replication within eukaryotic cells

To establish the respective role of the three Fis proteins in *Legionella* pathogenesis, single Δ*fis1*, Δ*fis2*, and double Δ*fis1*/Δ*fis2* (named hereafter Δ*fis1-2*) mutants of *L. pneumophila* Paris strain were constructed. Repeated attempts to obtain the Δ*fis3* mutant in either wild-type strain or single mutants failed, suggesting this gene is essential in this genetic context.

*Δfis1*, *Δfis2* and *Δfis1-2* mutants were assessed for their intracellular growth kinetics. Wild-type strain and mutants expressing the mCherry-fluorescent protein were used to infect either *Acantamoeba castellanii* amoebae or U937 macrophages, and the mCherry fluorescence, reflecting bacterial intracellular growth, was monitored over a period of 72h (**Fig. 2A and 2B**). As expected, fluorescence increases for wild-type strain but not for the avirulent *ΔdotB* mutant. Deleting *fis2* has minimal impact on virulence, while the *Δfis1* mutant is impaired in both eukaryotic hosts, showing a predominant role for Fis1 in intracellular replication. The *Δfis1-2* double mutant showed delayed and decreased intracellular growth, compared with the *Δfis1* mutant, revealing that Fis2 act synergistically with Fis1 to control intracellular replication of *L. pneumophila*. Overexpressing *fis1* or both *fis1* and *fis2* partially restores mutant phenotypes (**Fig. 2C**), confirming that Fis1 and, to a lesser extent, Fis2, play a role in the intracellular growth of *L. pneumophila*.

**Fig. 2.**
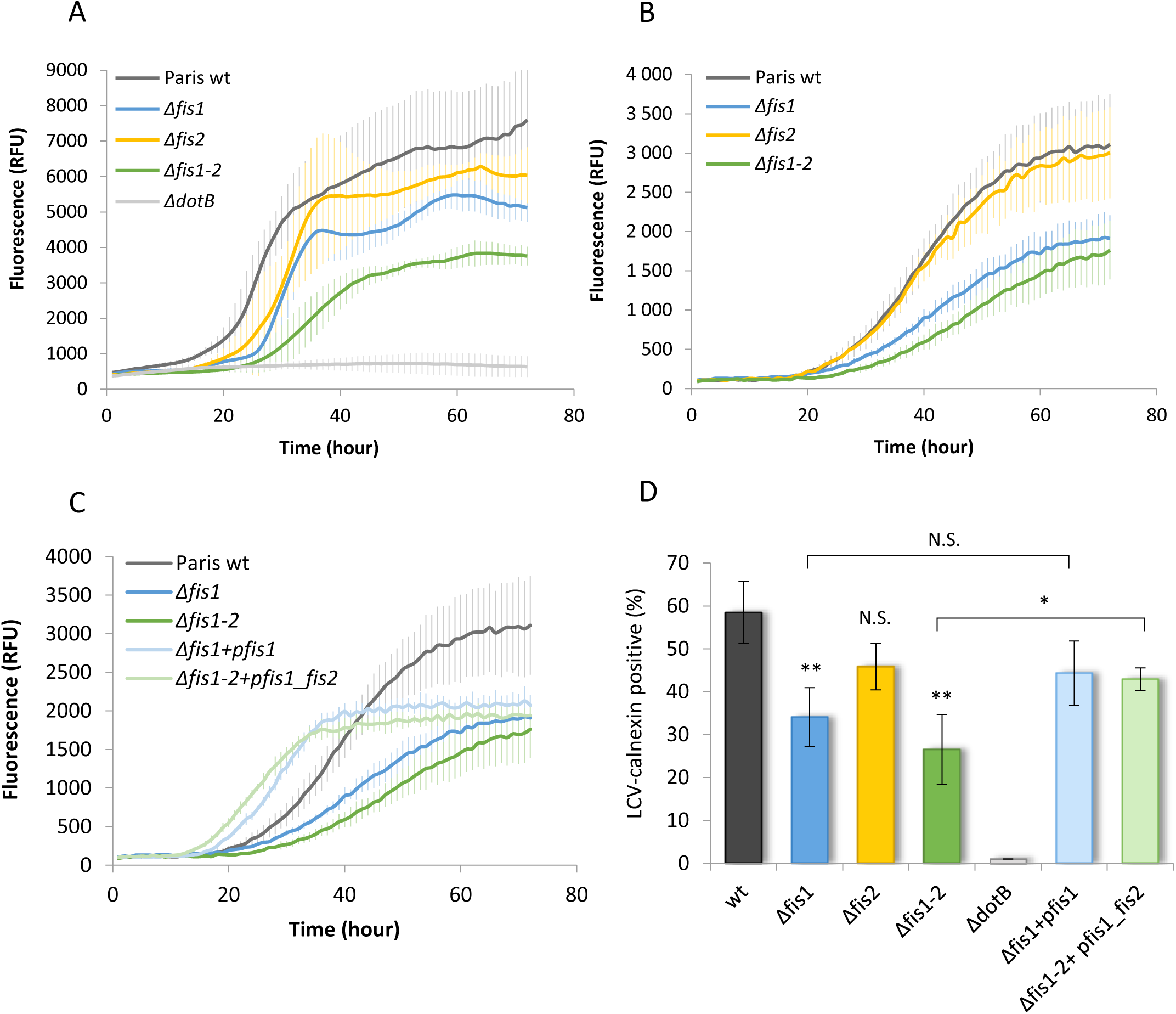
Intracellular growth and LCV biogenesis within eukaryotic cells depend on *L. pneumophila* Fis1 and Fis2 proteins. *Acanthamoeba polyphaga* (A) or U937 macrophages (B and C) were infected at an MOI of 10 with wild-type Paris strain (dark grey), Δ*fis1* (blue), Δ*fis2* (yellow), Δ*fis1-2* (green) and Δ*dotB* (light grey) mutants harbouring the mCherry-producing plasmid pXDC50. For complementation tests (C), Δ*fis1* and Δ*fis1-2* mutants carried pXDC50-derivative plasmids expressing either *fis1* gene (p*fis1*, hatched blue) or both, *fis1* and *fis2* genes (p*fis1_fis2*, hatched green), respectively (see Materials and Methods). Intracellular growth was monitored for 72 hours using a microtiter plate fluorescent reader and data, expressed in RFU (relative fluorescent units) represent the mean values with error bars obtained from experiments done in quadriplicate. The kinetics were performed at least three times and similar results were obtained. (D) LCV biogenesis is assessed by monitoring endoplasmic reticulum (ER) recruitment around the phagosome 1h post-infection. The amoeba *D. discoideum* producing GFP-calnexin fusion was infected at an MOI of 10 with mCherry expressing bacteria. For each strain, more than 200 vacuoles surrounded or not by green fluorescent ER-membrane were examined by confocal microscopy and the results are expressed in percentage of ER-recruiting LCVs (averages ± standard deviations of at least three independant experiments). Data were analysed with R software and significant differences were determined by using Tukey’s HSD test. *,** and N.S. indicate *P* <0.01, *P* <0.005 and non-significant, respectively.

Biogenesis of the endoplasmic reticulum (ER)-derived vacuole called LCV (*Legionella* Containing Vacuole), essential for *L. pneumophila* intracellular replication, was assessed by infecting GFP-calnexin producing *Dyctiostelium discoideum* amoeba with mCherry-expressing wild-type and mutant strains. Confocal microscopy showed that 57% of vacuoles with wild-type strain were ER-calnexin positive at 90 min post-infection **(Fig. 2D)**. As expected, the avirulent *ΔdotB* mutant failed to recruit ER vesicles. 47% of vacuoles with Δ*fis2* mutant were calnexin-positive, which is not significantly different from the wild-type, while only 36% and 27% of vacuoles with Δ*fis1* and Δ*fis1-2* mutants were calnexin-positive, indicating significant impairment in LCV biogenesis. Finally, ER-recruitment increases when the *Δfis1-2* mutant is complemented with a plasmid co-expressing both, *fis1* and *fis2* **(Fig. 2D)**.

Overall, these results confirm the key role of Fis1, but also the synergistic activity of Fis2, in controlling the virulence of *L. pneumophila*.

### Concomitant deletions of *fis1* and *fis2* have synergistic effects on global gene expression

To further characterize the respective role of Fis1 and Fis2 proteins in *L. pneumophila* virulence, we compared the global gene expression profiles of the *Δfis1, Δfis2* and *Δfis1-2* mutants. Importantly, we analyzed their transcriptome by RNA-Seq at the transition between the replicative and the transmissive phase of the infectious cycle (corresponding to early stationary phase broth cultures).

The deletion of *fis1* leads to significant expression changes, with 279 genes differentially expressed whereas the *Δfis2* mutant shows a less pronounced transcriptional impact, with only 59 gene expression changes compared to the wild-type strain (fold change (FC) >2) **(Fig. 3A)**. Notably, there is no overlap between the genes regulated by Fis2 and those controlled by Fis1, indicating that these 59 genes form a Fis2-specific regulon **(Fig. 3B)**. Intriguingly, the simultaneous deletion of both *fis1* and *fis2* results in a higher number of differentially expressed genes (609) than those observed in the *Δfis1* and *Δfis2* single mutants (279+59=338 genes). The genes exclusively identified in the double mutant might be targets of the combined action of Fis1 and Fis2 or might have been undetected in the single mutants due to an FC below the threshold of 2. Consistently, 60% of the genes targeted by Fis1 show a greater FC in the double *Δfis1-2* mutant. These data are consistent with the respective virulence phenotypes of the *Δfis1* and *Δfis2* mutants described above. They highlight the predominant role of Fis1 and show that Fis1 and Fis2 share common target genes they synergistically regulate.

**Fig. 3.**
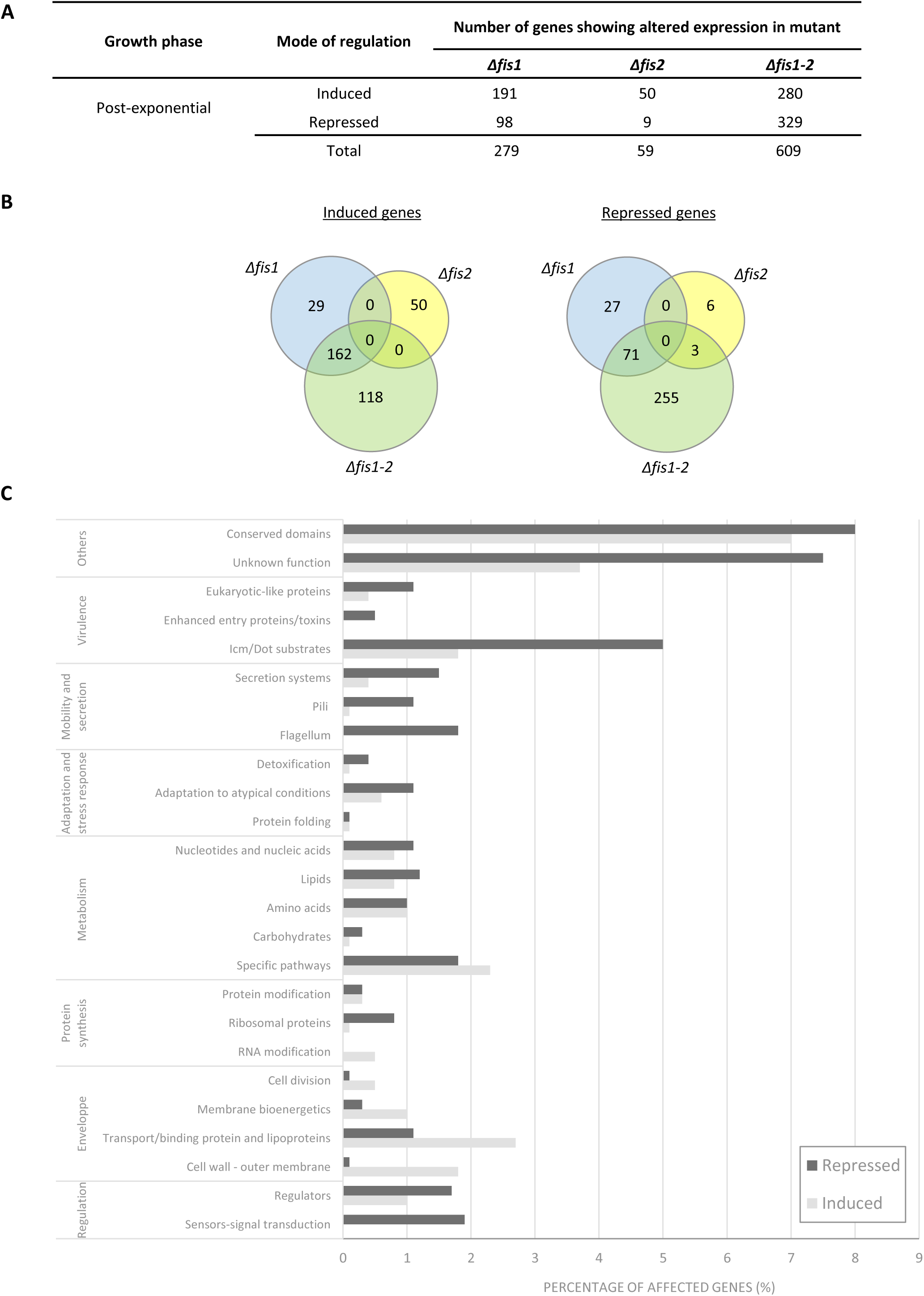
Whole-transcriptome analysis obtained from total RNA sequencing (RNA-Seq) of *Δfis1, Δfis2* and *Δfis1-2* mutants compared to wild type strain. (A) The number of genes positively or negatively regulated in post-exponential (PE) phase is indicated for each mutant (the count excludes genes located on plasmids). (B) Venn diagrams show the total number of differentially expressed genes (DEGs) in absence of Fis1, Fis2 or both proteins (0,5>FC>2 and FDR-corrected *P*<0,05). (C) The genes positively or negatively affected in the *Δfis1-2* double mutant were classified into 10 arbitral main functions and further divided into functional categories (y axis) according to their annotation or prediction defined in the *L. pneumophila* genome Paris (Cazalet et al. 2008). The percentage in each subcategory corresponds to the number of genes upregulated (or downregulated) in this category out of the total number of affected genes x100.

The virulence defects of the *Δfis1* and *Δfis1-2* mutants are reflected in their global gene expression profiles. Many transmissive genes such as flagellar and type IV pili genes or several effector-encoding genes (51) are downregulated in mutants (**Fig. 3C**). T4SS-independent virulence factors like HtpB (Hsp60), LadC, RtxA toxin, as well as the enhanced entry proteins EnhABC and their homologues are also repressed in Δ*fis1-2* mutant. Moreover, key post-exponential regulators are modulated in a Fis1/Fis2-dependent manner. The transmission trait enhancer LetE, the RNA chaperone Hfq, sigma factors RpoN (σ54), RpoE (σ24), and FliA (σ28), along with TCS-response regulators PilR, CpxR and PmrA, are all down-regulated in the *Δfis1-2* mutant strain (**Table S2**). In addition, 9 of 24 GGDEF/EAL regulatory proteins which play a key role in the control of virulence, biofilm formation and flagellar regulation (69–72) show lower expression in the *Δfis1-2* mutant. Overall, these data suggest that Fis1 and Fis2 are part of the regulatory circuit that governs the biphasic life cycle of *Legionella*.

### Fis1 and Fis2 synergistically control the expression of flagellum and type IV pili

To further investigate the role of Fis1 and Fis2 in controlling the biphasic infectious cycle of *L. pneumophila*, we deciphered in detail their role in the expression of two important transmissive traits associated with *Legionella* virulence, namely the flagellum and type IV pili.

The biogenesis of functional flagella involves 45 flagellar genes organized into four distinct classes, controlled by a hierarchical cascade initiated by the major regulators FleQ and RpoN (87). Remarkably, 24 flagellar genes, including RpoN and FliA sigma factors required for the transcription of flagellar genes, are downregulated in the *Δfis1-2* double mutant (**Fig. 4A**). To confirm that Fis1 and Fis2 control flagella biogenesis, the production of the flagellin FlaA was quantified by immunoassay using an anti-FlaA antibody (**Fig. 4B**). Flagellin is not detected in exponential phase but synthesized by wild-type strain, *Δfis1* and Δ*fis2* in post-exponential phase. More importantly, FlaA synthesis is strongly decreased in *Δ*fis*1-2*, similar to non-flagellated *ΔfleR* mutant (73), showing that Fis1 and Fis2 cooperate to control synthesis of flagellin in *L. pneumophila*. Overexpressing *fis1* or *fis2* in *Δfis1-2* mutant restores flagellin production, but Fis3 does not complement this phenotype suggesting a distinct regulatory function. Consistent with immunoassay results, electron transmission microscopy observations show that the wild-type strain, *Δfis1* and *Δfis2*, but not the *Δfis1-2* double mutant, display flagella (**Fig. 4C**). Together, these data demonstrate that Fis1 and Fis2 jointly control the expression of flagellar genes and the biogenesis of the flagellum.

**Fig. 4.**
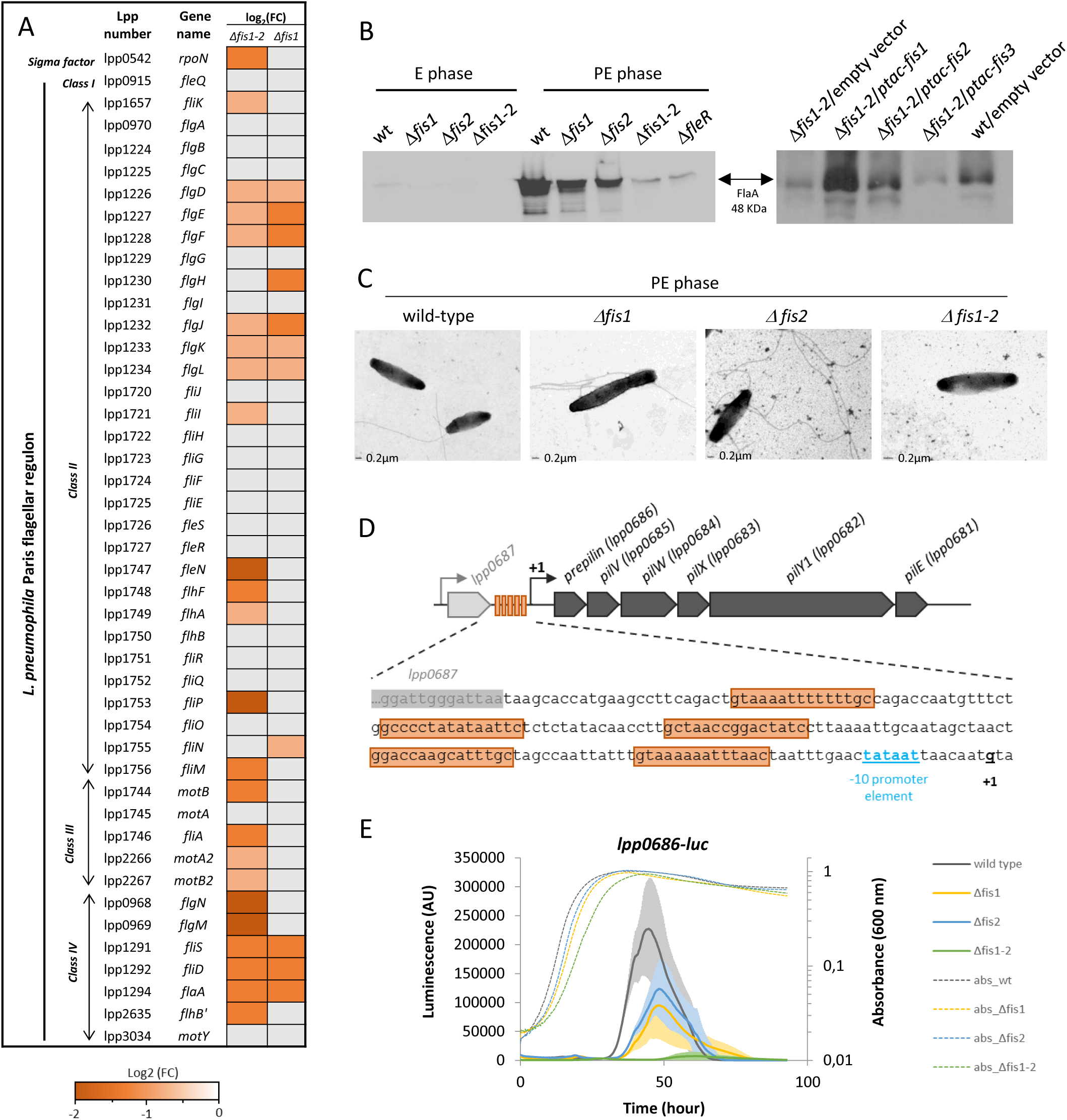
Effect of *fis* genes deletion on flagellum biosynthesis and pilus gene expression in *L. pneumophila* Paris. (A) Flagellar genes significantly downregulated in the Δ*fis1* and Δ*fis1-2* mutants (FDR-corrected *P* ≤ 0,05). Their expression pattern (expressed by log2FC) is depicted in a color code based on the color scale given below. Light grey box means that the expression does not change or is not significant. (B) Western blot analysis of whole-cell lysate of wild type strain, Δ*fis* mutants and *fleR* mutant strain (negative control), probed with a FlaA polyclonal antibody (left panel). Cells were harvested during exponential (E) or post-exponential (PE) growth phase.The right panel showed the expression of flagellar FlaA protein in PE-phase cell lysates of the Δ*fis1-2* double mutant complemented with either *fis1*, *fis2* or *fis3* genes and the Δf*is1-2* and wild-type strain harbouring empty vector. (C) Transmission electron microscopy pictures of post-exponential (PE)-phase *L. pneumophila* Paris wild-type, Δ*fis1*, Δ*fis2* and Δ*fis1-2* mutants. Flagellar structure is absent only in the Δ*fis1-2* double mutant strain. (D) Genomic organization of pilus-encoding operon and the DNA sequence located upstream the operon. The -10 promoter element is in italic, the transcription start is in boldface and underlined and the putative Fis-regulatory elements are shaded in orange. (E). The transcription activity of the pilus-related operon (*lpp0686*-*lpp0681*) is examined in the wild-type (orange), Δ*fis1* (yellow), Δ*fis2* (blue) and Δ*fis1-2* (green) mutants by the means of the *lpp0686*-*luc* transcriptional fusion. Luciferase activity (solid lines) and absorbance (dotted lines) were monitored for 96 hours as described in the Experimental Procedures. Data expressed in Relative Light Units (RLU) are means and standard deviations of technical triplicate and representative of at least three independant experiments.

Type IV pili are also considered an important transmissive trait (50, 51). The operon *pilBCD*, the type IV prepilin gene *pilE1*, the major type IV pilin *pilE2* and the operon clustering *pilE3* (*lpp0686* to *lpp0681*) are all down-regulated in the *Δfis1-2* double mutant compared to the wild-type strain (**Table S2**). To confirm the RNAseq data, a transcriptional fusion containing the 300-bp promoter region of the pilus operon *lpp0686-lpp0681* fused to the firefly luciferase gene *luc* (designated P*pilE-luc*) was constructed in the wild-type strain and mutants. Noteworthy, 4 putative Fis binding sites were identified in this regulatory region, based on similarities with the 15-bp consensus sequence established for *E. coli* (74, 75) (**Fig. 4D**). Maximum luminescence was observed after bacteria enter the stationary phase, confirming that pili are transmissive traits. Importantly, P*pilE-luc* expression was repressed in *Δfis1* and *Δfis2* mutants and nearly abolished in *Δfis1-2* double mutant, indicating that pilus gene expression depends on both Fis1 and Fis2 (**Fig. 4E**).

Taken together, these results strongly support a significant role of Fis1 and Fis2 in the regulation of key virulence factors known to be induced during the transmissive phase.

### Fis1 and Fis2 are essential for optimal Dot/Icm-dependent translocation

To better understand the roles of Fis1 and Fis2 in *L. pneumophila* virulence, we examined their impact on the regulation of T4SS Dot/Icm synthesis which secretes over 300 effector proteins into the host cell and is crucial for intracellular replication. Among the 27 *dot/icm* genes organized into 14 transcription units (92) **(Fig. 5A)**, *icmPO* (*dotML*), *icmDJB* (*dotPNO*), *icmV-dotA*, *icmWX* operons, as well as the *dotV* gene are downregulated, whereas the *dotKJIHGF* operon is upregulated in the *Δfis1-2* double mutant. Moreover, several putative *fis* binding motifs are identified in the promoter regions of these transcriptional units (**Fig. 5A**). To analyse the effect of Fis1 and Fis2 on Dot/Icm synthesis, we examined the expression of four transcriptional *luc*-fusions with the promoter regions of gene or operon (i) *icmTS* (designated P*icmT*-*luc*), (ii) *dotKJIHGF* (P*dotK*-*luc*), (iii) *dotV* (P*dotV*-*luc*) and iv) *icmV*-*dotA* (*PicmV-luc*), during axenic growth. The expression of P*dotV*-*luc* and P*icmV*-*luc* decreased, while P*dotK*-*luc* increased in the *Δfis1-2* double mutant compared to the wild-type strain **(Fig. 5B)**, confirming the RNAseq data and the synergistic effect of Fis1 and Fis2. Interestingly, P*icmT-luc* expression was 2-fold higher in all three *fis* mutants, which was not apparent in the RNAseq data **(Fig. 5A and 5B)**. This discrepancy is likely because P*icmT-luc* expression differences between Δ*fis1-2* and wild-type strain are most pronounced during the exponential growth phase, while RNAseq was performed on post-exponential phase cultures. These results demonstrate that Fis proteins negatively regulate the *icmTS* operon, likely through binding to the putative regulatory regions identified.

**Fig. 5.**
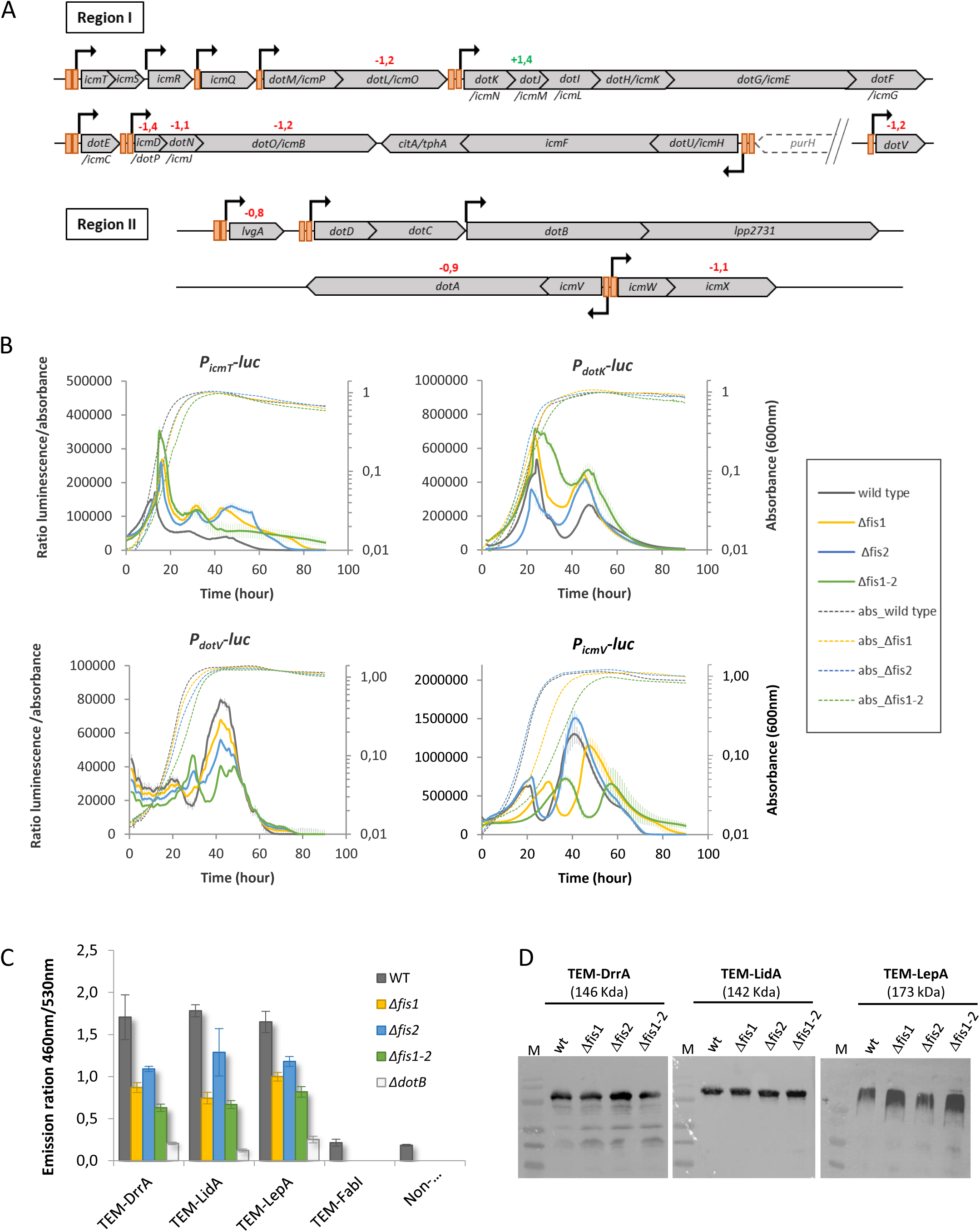
The absence of Fis1 and Fis2 impact *icm*/*dot* genes expression and the functioning of the T4SS apparatus in *L. pneumophila*. (A) The *icm/dot* genes are distributed in two distinct genomic regions on the chromosome and organized in several operon structures whose promoters are indicated by black arrows. Orange boxes located in these promoter regions indicate the presence of putative Fis binding sites (2 boxes mean the presence of 2 or more sites). The significant fold-change (FC) expression data of the Δ*fis1-2* mutant *vs.* wild-type strain is indicated above the corresponding gene, in red for repressed genes and green for induced ones. (B) The expression level of transcriptional fusions *PicmT-luc (*operon *icmTS), PdotK-luc (*operon *dotKJIHGF*), *PdotV-luc and PicmV-luc* (operon *icmV*-*dotA)* were monitored for 90 hours in wild-type strain (dark grey), Δ*fis1* mutant (blue), Δ*fis2* mutant (yellow) and Δ*fis1-2* double mutant (green). Solid lines and dotted lines represent luciferase activity and absorbance, respectively. (C) The translocation efficiency of three distinct β-lactam TEM-effector fusions (TEM-drrA, TEM-lidA, TEM-lepA) was monitored in the wild-type strain and the Δ*fis1,* Δ*fis2,* Δ*fis1-2,* Δ*dotA* mutant strains. The TEM-fabI fusion, coding for a cytoplasmic protein, transformed in the wild-type strain was used as a negative control of translocation. Differentiated-U937 cells were infected with strains at a MOI of 10 and the infection was allowed to proceed for 1 h before β-lactam substrate (CCF4-AM) was added. Translocation efficiency is calculated as the ratio of cleaved and uncleaved CCF4. (D) The expression level of the three TEM effector fusions in the different genetic backgrounds was analyzed by immunoblot using an anti-TEM antibody to confirm a similar expression level in all strains.

Since Fis1 and Fis2 controlled *dot/icm* gene expression, we tested T4SS functionality in Δ*fis* mutants, by measuring translocation of several Dot/Icm effectors (LepA, DrrA, LidA) using the β-lactamase reporter (71, 76). U937 phagocytes were infected with Paris strain, *ΔdotB*, or *Δfis* mutants expressing TEM-effector fusions, and effector secretion was quantified by CCF4 fluorescence change from green to blue. As expected, TEM-LepA, TEM-DrrA, and TEM-LidA were efficiently secreted by the wild-type strain (460nm/530nm ratio >1.5), but not by *ΔdotB*. Translocation decreased in *Δfis1* (50%), *Δfis2* (45%), and *Δfis1-2* (65%) mutants (**Fig. 5C**). Importantly, TEM fusion proteins are produced in similar and stable amounts across different genetic backgrounds **(Fig. 5D)**, indicating that the reduced TEM fusion protein detected in the host cell is due to lower translocation efficiency by the Dot/Icm machinery, rather than reduced synthesis by the bacteria. These data confirm Fis1 and Fis2 impact both *dot/icm* gene expression and T4SS functionality, thus contributing to *Legionella* virulence.

### Fis proteins positively or negatively regulate the expression of numerous T4SS effectors

The Dot/Icm T4SS secretes over 300 effector proteins into the host cell during infection, with secretion controlled over time to match their biological function. This temporal regulation is achieved by controlling the Dot/Icm machinery’s functionality (69) and effector gene expression. Most effectors are synthesized in the post-exponential growth phase, similar to the transmission phase of the *Legionella* infection cycle, though some are produced during the exponential phase (77, 78).

Transcriptomics of *Δfis1*, *Δfis2*, and *Δfis1-2* mutants identified 17 upregulated and 44 downregulated effector genes (fold-change > 1.5) **(Table 1)**. Many genes altered in *Δfis1* are more affected in *Δfis1-2*, showing Fis1 and Fis2 synergistic action. Among the 17 effector genes repressed by Fis1 and Fis2, *ankC*, *legK3* and *cegK3* were previously identified as being inhibited by Fis1 and Fis3 in the Philadelphia strain of *L. pneumophila* (26, 67). Conversely, the *sidC* gene, strongly repressed by Fis3 in Philadelphia (26), appears to be activated by Fis1 and Fis2 in Paris strain. Among the 44 effector genes activated by Fis1 and Fis2, 29 have been identified as typical virulence factors of the transmissive phase (51, 78). Interestingly, this group includes 11 effectors known to be repressed by the PmrA/B TCS (60, 61), excepted *sidG* which is activated by PmrA and *sidH* whose expression is enhanced by both PmrA and CpxR. Additionally, *ralF*, *sidC* and *sdcA* are also under the positive regulation of the *Legionella* quorum sensing (Lqs) system (57, 58). These data suggest that effector gene promoters are subject to multiple transcriptional controls, including the three Fis proteins, to fine-tune the synthesis of each effector during the infectious cycle.

To experimentally confirm the activating or repressive regulatory role of Fis1 and Fis2, we selected 4 differently regulated effector genes, namely RalF, SidG, AnkG, and MarB to analyze the expression of the related transcriptional *luc*-fusions in the Paris wild-type strain, *Δfis1*, *Δfis2*, and *Δfis1-2* mutants over 90h of growth in axenic medium. Expression profiles of these fusions confirm the RNAseq data, showing that 2 genes are positively controlled by Fis1 and Fis2 (*ralF* and *sidG*, **Fig. 6A**) and 2 genes are negatively controlled (*ankG* and *marB,* **Fig. 6B**). Interestingly, the positively regulated genes are strongly expressed in post-exponential phase while the negatively regulated genes are mainly expressed during exponential phase, suggesting a growth phase-dependent regulatory role of Fis. Notably, the absence of Fis2 alone affects the expression levels of P*ralF-luc* and P*sidG-luc*. This effect was not detected in RNAseq, likely due to the higher sensitivity of *luc*-reporter fusions compared to the global transcriptomic approach. Overall, these results highlight the regulatory function of Fis1 and, for the first time, uncover the role of Fis2 in effector gene expression. Additionally, they reveal that the repertoire of effector genes regulated by Fis proteins is broader than previously described (26).

**Fig. 6.**
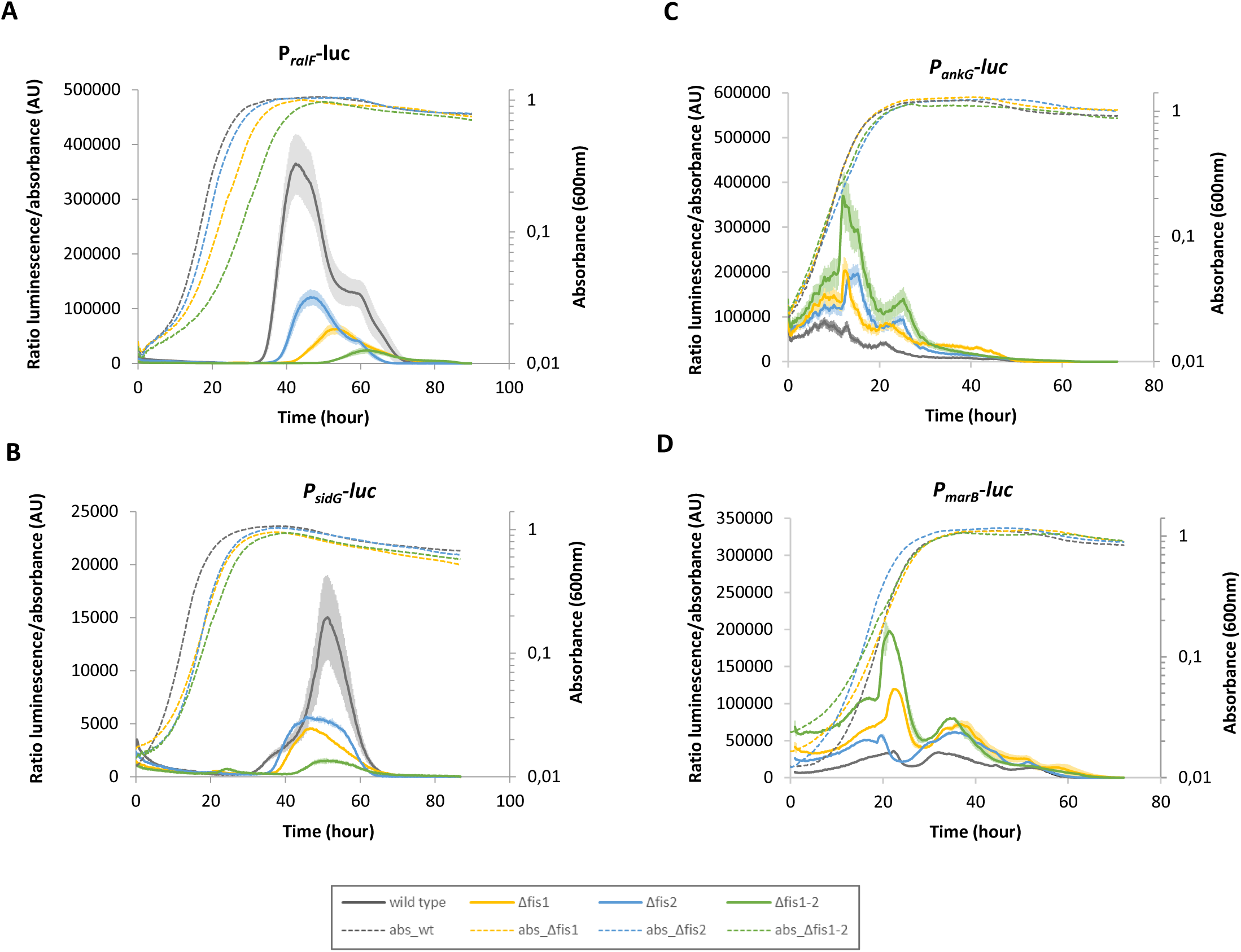
Different effector genes are regulated by Fis proteins either positively (A and B) or negatively (C and D). The expression level of *luc* fusions of four effector-encoding genes (*ralF*, *sidG*, *ankG and marB*) was examined in wild-type strain (dark grey), Δ*fis1* mutant (blue), Δ*fis2* mutant (yellow) and Δ*fis1-2* double mutant (green). Luciferase activity (solid lines) and absorbance (dotted lines) were monitored for 96 hours as described in the Experimental Procedures. Data expressed in Relative Light Units (RLU) are means and standard deviations of technical triplicate and representative of at least three independant experiments.

### Fis1 and Fis2 bind Fis-consensus regulatory elements in the promoter of their targeted genes

As previously noted, we identified putative Fis-binding sites in the promoters of genes targeted by Fis1 and Fis2 proteins. Positions 1 (G) and 15 (C) of these sites, indicated by asterisks in **Fig. 7A**, were substituted using site-directed mutagenesis and the resulting mutated promoter regions were fused to the luciferase gene, creating PpilE_mut-*luc*, P*ralF_mut*-*luc* and P*icmT_mut*-*luc* fusions. Mutations introduced into Fis consensus binding sites of *ralF* and *pilE* fusions result in a sharp decrease of luciferase, and conversely in a significant increase of expression of the P*icmT_mut*-*luc* fusion, thus confirming the direct regulation of the 3 genes by Fis1 and Fis2 proteins, likely through Fis binding to the sites identified in their promoters (**Fig. 7B**). To confirm this binding, Electrophoresis Mobility Shift Assays (EMSA) were performed using native purified Fis1 and Fis2 proteins and PCR fragments encompassing the putative Fis regulatory elements of each promoter as binding targets. As the amount of Fis1 or Fis2 increased, shifted DNA bands were observed, indicating that both Fis1 and Fis2 can bind to the *pilE*, *ralF*, and *icmT* promoters **(Fig. 7C)**. Notably, at equal protein concentration (100 µM), all the DNA molecules are shifted with Fis1 but not with Fis2, demonstrating that Fis1 binds the promoters with greater affinity than Fis2, which is consistent with the lesser impact of Fis2 on the expression of these genes, as observed by RNAseq and *luc*-fusions. Moreover, while only a single high-molecular-weight shifted band is observed for Fis1, indicating its simultaneous binding to multiple sites, distinct lower molecular weight shifted bands appear as the Fis2 concentration increases. These bands correspond to different Fis2-DNA complexes, suggesting that Fis2 binds to fewer sites and in a more gradual manner compared to Fis1. Finally, the specificity of Fis1 and Fis2 binding to the *lpp0686*, *ralF*, and *icmT* promoters was demonstrated, as both proteins bind with much less efficiency to the mutated target DNA **(Fig. 7C)**. In the cases of the mutated-*pilE* and -*ralF* promoters, low-intensity shifted bands were observed, suggesting that Fis1 and Fis2 can still bind to other putative sites in these promoter regions **(Fig. 7A)**.

**Fig. 7.**
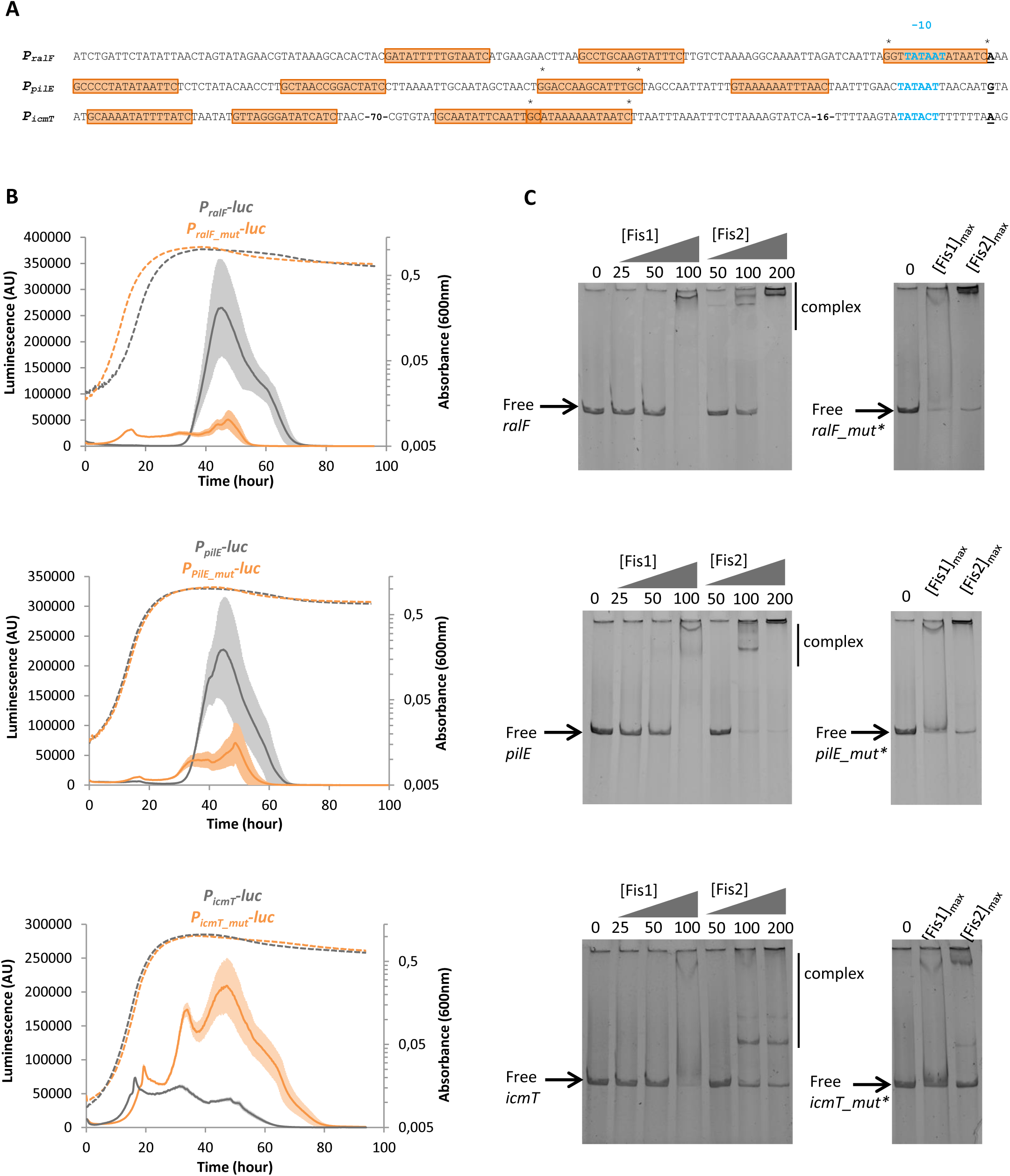
*L. pneumophila* Fis1 and Fis2 proteins bind and regulate the promoter regions of *ralF*, *pilE* and *icmT* genes. (A)The regulatory regions of *ralF*, *pilE* and *icmT* with their putative Fis-binding sites (in orange boxes), the -10 promoter element (in blue) and transcription start (bold, underlined) are represented. The mutations introduced in the putative Fis site are marked by asterisks (B) The expression level of wild-type *ralF-luc*, *pilE-luc* and *icmT-*luc-fusions (in grey) and the same fusions with mutations in the putative Fis-binding sites (*ralF_mut-luc*, *pilE_mut-luc* and *icmT_mut-*luc, in orange) were examined in the wild-type strain genetic context. (C) Electrophoretic mobility shift assays were performed with purified PCR-amplified DNA fragment of promoter regions and increasing concentrations of purified native proteins Fis1 (0, 25, 50, 100 µM) and Fis2 (0, 50, 100, 200 µM) (left panel). On the right panel, the binding experiments were performed by incubating the DNA fragment containing mutations in the Fis binding site with the highest protein amount used in the left panel. The first lane of each panel did not contain any protein (free DNA).

### Fis1 and Fis2 display redundant activity and partially distinct growth-dependent expression pattern

We sought to investigate whether the cooperation of Fis1 and Fis2 in controlling virulence traits is due to their complementarity, interchangeability, or more complex interactions. To address the question of the role of *Legionella* Fis proteins in a simple way, we tested their ability to complement phenotypes characteristic of an *E. coli Δfis* mutant, namely (i) the lack of mobility due to the absence of activation of flagellum synthesis (79), and (ii) a dysregulation of DNA supercoiling due to the absence of repression of the DNA gyrase gene (80, 81)(**Fig. 8A**). We also assayed each *Legionella* Fis protein to control the expression of the *E. coli fis* gene, i.e. to reproduce the autoregulation set up by the *E. coli* Fis protein (82). We first deleted the *fis* gene in *E. coli* BW25113 to obtain the *Δfis* mutant strain BWL0, and then replaced the *E. coli fis* gene by *fis1, fis2* or *fis3* from *L. pneumophila* under the *E. coli fis* promoter (**Fig. 8B**). Noteworthy, we were unable to obtain *E. coli* strain expressing *fis3* gene, as genetic reconstruction led to mutations in the 3’ end of *fis3* gene. As expected, deletion of *E. coli fis* results in both reduced motility (**Fig. 8C**), increased expression of *fis* (**Fig. 8D**), and activation of P*gyrA* (**Fig. 8E**), when compared to the *E. coli* wild-type strain. All 3 phenotypes are complemented by the expression of *Legionella fis1* or *fis2* genes, with Fis1 being better able to complement all phenotypes. This suggests that Fis1, and to a lesser extent Fis2, have similar activities to *E. coli* Fis. Better complementation by Fis1 could be consistent with its higher activity and its predominant role in the control of phase transition and virulence of *L. pneumophila*. Furthermore, the inability to express *fis3* in *E. coli* likely highlights a different role for Fis3.

**Fig. 8.**
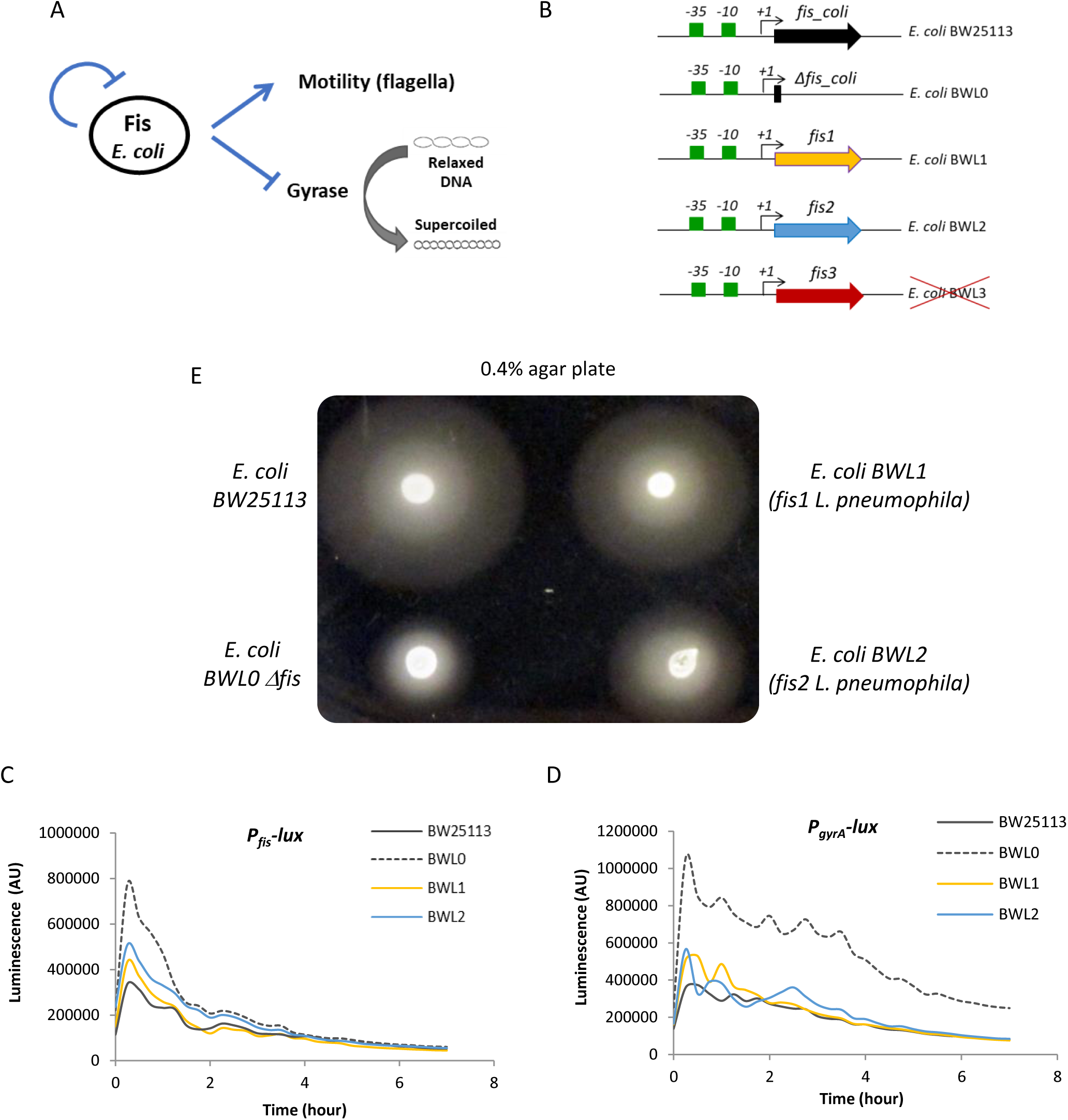
Ectopic expression of *Legionella* Fis1, Fis2 and Fis3 in *E. coli* BW25113. (A) Schematic representation of regulation of motility (flagella), DNA topology (gyrA) and *fis* gene expression by Fis in *E. coli*. Arrows and blunt arrows represent positive or negative regulation, respectively. (B) Schematic representation of the different *E. coli* strain: *E. coli* BW25113 is used as the wild-type strain; BWL0 is a *fis* null mutant and BWL1 et BWL2 express *fis1* and *fis2* gene, respectively instead of the endogenous *fis* gene. It was not possible to construct the BWL3 strain expressing *fis3* of *L. pneumophila*. (C) Swimming motility of *E. coli* strains on 0,3% soft agar plate. Bacteria were stabbed on the agar plate and incubated at 30°C for 16h. A positive motility was indicated by diffused growth around the point of inoculation while growth restricted around the stab point indicated a non-motile strain. (D and E) The expression level of the transcriptional fusions Pfis-lux and PgyrA-lux were examined in the *E. coli* BW25113 wild type strain (black), BWL0 Δ*fis* mutant (dashed black), BWL1 mutant (yellow) and BWL2 mutant (blue). Luciferase activity was monitored for 6 hours.

As Fis1 and Fis2 have redundant functions and target the same DNA regions in *L. pneumophila*, we aimed to determine the relative abundance of the 3 Fis proteins throughout axenic growth. We analyzed samples extracted at OD600 = 1; 2; 3; 4 and 5 from 3 independent liquid growth of Paris strain at 30°C using quantitative mass spectrometry **(Fig. 9)**. Fis1 and Fis3 are highly expressed early in growth (OD600 = 1), then decrease to a minimum in the stationary phase (OD600 = 5). Conversely, Fis2 levels are low initially, peak at the end of the exponential growth phase (OD600 = 3), and then decrease. In Philadelphia strain grown at 37°C Fis1-Fis3 are highly, and Fis2 exclusively, expressed during the exponential phase (77), similarly to the unique Fis protein in *E. coli* and other enteric bacteria that peaks at the beginning of the logarithmic growth phase (23, 83–85).

**Fig. 9.**
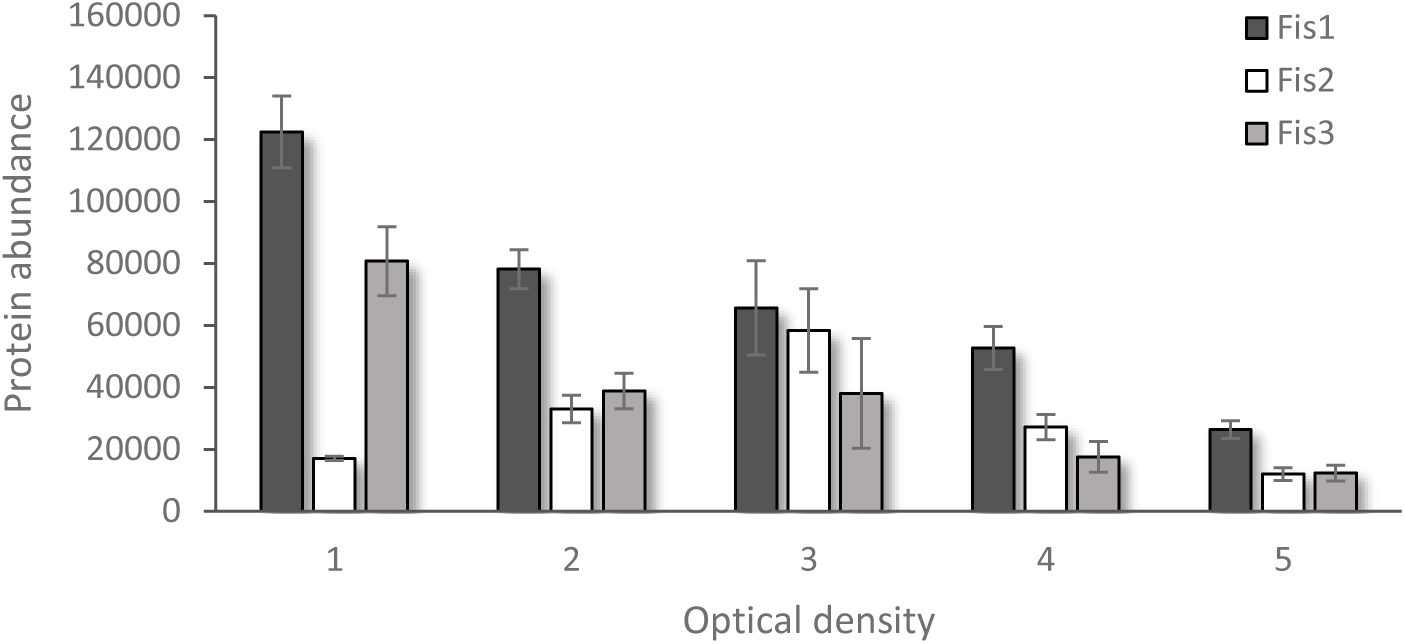
Expression level of Fis1, Fis2 and Fis3 during bacterial growth. Three independent cultures of *L. pneumophila* Paris WT were grown in liquid medium AYE (at 30°C). At the desired OD600 (1, 2, 3, 4 and 5), samples were collected, and their protein content was analyzed by mass spectrometry after sample-specific labelling (see Material and methods for details). The graph represents the mean relative abundance of each Fis proteins with the standard error of the mean. Complete data set is presented in Table S3.

The dynamic presence and absence of the 3 *Legionella* Fis proteins likely reflect distinct and specific regulatory roles. Noteworthy, despite their disparate production profiles, Fis proteins coexist at specific points in time and are likely to collaborate during the *Legionella* infection cycle.

### Fis1 and Fis2 could form heterodimers to cooperatively control *L. pneumophila* virulence

From a biochemical perspective, Fis1, Fis2, and Fis3 proteins of *L. pneumophila* have limited amino acid sequence similarities with each other and with the canonical Fis protein of *E. coli* or *Salmonella*, however, they show remarkable structural similarities **(Fig. 10A)**. According to AlphaFold models, their quaternary structure as homodimers is similar. The α-helices from each subunit (α1-α2 and α1’-α2’) assemble into a four-helix bundle forming the Fis dimer core. Additionally, the helix α3-α4 and α3’-α4’ at the C-terminal end forming a helix-turn-helix (HTH) motif involved in DNA recognition and binding (26) is also well conserved. Conversely, the N-terminal end exhibits greater variability. Fis1 and Fis2 display a highly flexible end while the N-terminus of Fis3 forms a continuous helix with helix α1, both different from the N-terminus of Fis of *E.coli* which folds into 2 β-hairpins required for controlling Hin-catalyzed DNA inversion (86–88) **(Fig. 10B)**. Importantly, critical residues for DNA invertases activation in *E. coli*, are not conserved in *Legionella* Fis proteins, which raises questions about the role of *Legionella* Fis N-terminus in DNA inversion regulation or other DNA transactions. Interestingly, AlphaFold modelling also yielded very reliable predictions of Fis1-Fis2, Fis1-Fis3, Fis2-Fis3 heterodimeric complexes (**Fig. 10B**), supported by pLDDT and idTT scores similar to the homodimer *E. coli* Fis model.

**Fig. 10.**
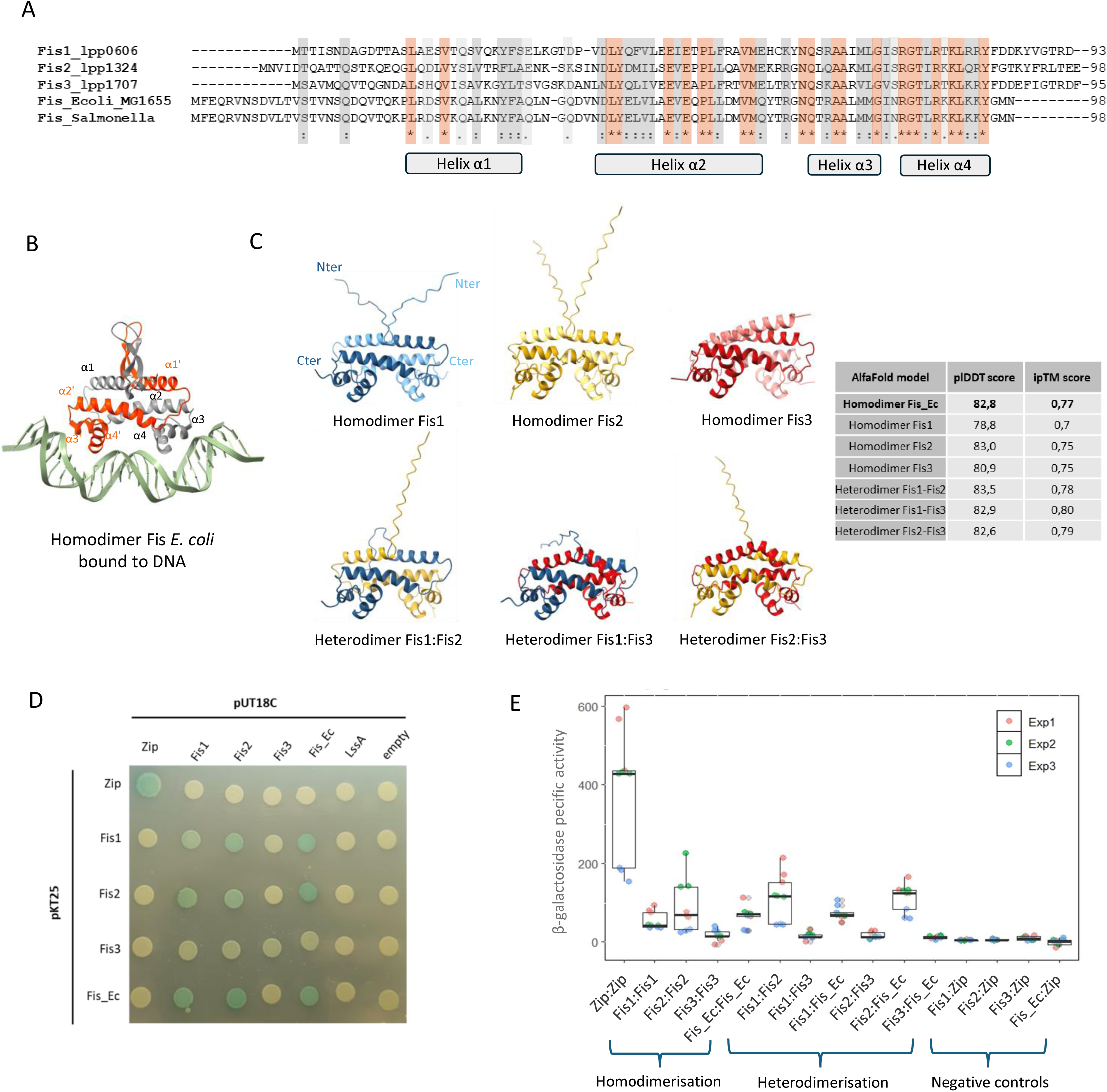
*Legionella* Fis proteins can form homodimers or heterodimers. (A) Sequence alignment of Fis proteins from different bacteria: the three Fis proteins of *L. pneumophila* Paris, Fis from *E. coli* MG1665 and *Salmonella enterica* serotype Thyphimurium (B) Crystal structure of Fis *from E. coli (*Fis_Ec) bound to 27 bp optimal binding sequence F1 (PDB doi: https://doi.org/10.2210/pdb3IV5/pdb). (C) AlphaFold models of *L. pneumophila* Fis1, Fis2, Fis3 homo- and heterodimer were generated by AlphaFold Multimer (ColabFold/AlphaFold2.ipynb, Mirdita *et al.* 2022). The per-residue confidence (pLDDT) metric and the interface predicted template modelling (ipTM) score were given for evaluating the accuracy of the predictions. (D) Representative image of qualitative β-galactosidase assays on agar plate. Overnight cultures of *E. coli* BTH101 co-expressing pKT25 and pUT18C derivatives were spotted on LB-X-Gal plates, which were then incubated for at least 48 h at 30 °C. pKT25 fusions are shown in column and pUT18c fusions in rows, with the indicated protein, empty vector or Zip positive control. The protein LssA encoded by the *lssXYZABD* locus which constitute the core components of the *Legionella* T1SS is used as a negative control as it is not expected to interact with Fis proteins. (E) The bacterial adenylate cyclase two hybrid (BACTH) system was used to study protein–protein interactions between Fis1, Fis2, Fis3 and the Fis protein of *E. coli* (Fis_Ec). Specific β-galactosidase activity was measured as described in Materials and Methods. Data represent the averages ± standard deviations of three technical replicates and three independent experiments. Statistical tests were performed between the different conditions (Figure S3).

To confirm that *Legionella* Fis proteins can form homo- and heterodimers, we performed bacterial two-hybrid assays between Fis1, Fis2, Fis3, and *E. coli* Fis. Briefly, Fis proteins were fused to the T25 and T18 fragments of the adenylate cyclase (CyaA) catalytic domain. *E. coli cya* mutant BTH101 was co-transformed with plasmids bearing T25 and T18 fusion proteins. Interaction between hybrid proteins restores CyaA enzyme function, leading to cAMP synthesis, which activates the *lac* operon via the cAMP-CAP complex. Protein-protein interaction was thus assessed by measuring β-galactosidase activity on X-Gal plates **(Fig. 10D)** and in ONPG - liquid assays **(Fig. 10E)**. *E. coli* Fis, and *Legionella* Fis1 and Fis2 can form homodimers, as the corresponding strains form blue colonies on X-Gal and significant β-galactosidase activity is detected in ONPG - liquid assays. Conversely, Fis3 transformed strain displays white/pale blue colonies and low β-galactosidase activity, indicating it does not form homodimers or the complex is unstable. More interestingly, Fis1 and Fis2 are able to form heterodimers with each other and with *E. coli* Fis. These interactions are specific, as shown by negative controls using LssA, a Type 1 secretion complex protein, and the Zip segment **(Fig. 10D and Figure S3)**. It is tempting to speculate that Fis1 and Fis2 heterodimers might recognize regulatory sequences with different affinities compared to homodimers. Given the varying abundance of Fis1 and Fis2 during the *Legionella* infection cycle, it can be postulated that the proportion of hetero- and homodimers is likely to vary during intracellular growth. This introduces a further layer of complexity and subtlety to the regulation of Fis.

## DISCUSSION

The virulence of pathogenic bacteria is based on complex strategies involving the finely regulated expression of different virulence genes. Appropriate control of this expression is achieved by sophisticated regulatory networks at the transcriptional, post-transcriptional, translational and post-translational levels. These targeted controls, specific to each virulence gene, are complemented by the action of global regulators, including NAPs, whose role in virulence has been somewhat neglected. Our study contributes to this field by demonstrating the role of Fis1 and Fis2 in controlling *L. pneumophila* virulence. Fis proteins have already been shown to participate to the virulence of several human pathogens such as *E. coli* pathogenic strains (18, 19), *Shigella flexneri* (18), *Haemophilus ducreyi* (20), *Yersinia pseudotuberculosis* (21), *Pseudomonas aeruginosa* (17) and its role been extensively studied in the human pathogen *Salmonella enterica* where it regulates the sequential expression of the *Salmonella* pathogenicity islands SPI-1 and SPI-2, both of which encode type 3 secretion systems (T3SS) and effector proteins required for invasion of host tissues or macrophages (23, 24, 89). The significance of our discovery lies in the fact that *L. pneumophila* has three non-homologous Fis proteins instead of just one, making it a unique model for dissecting the role of the Fis network in virulence. Specifically, we have shown that deletion of *fis1* has a major impact on *L. pneumophila* virulence, leading to defects in LCV biogenesis and subsequent intracellular replication, and that deletion of *fis2* increases the intensity of these phenotypes. Consistently, Fis1 plays a predominant role in gene regulation, as the transcriptome of the *Δfis1* mutant shows 279 deregulated genes, the *Δfis2* mutant only 59, and the *Δfis1-2* double mutant more than 600 genes. Noteworthy, we were unable to dissect the role of Fis3 in the virulence of *L. pneumophila* Paris due to our inability to delete the *fis3* gene. This suggests that *fis3* is essential in the genetic background of the Paris strain, whereas it is not essential in the Philadelphia background (26). Consistent with our results, it is interesting to note that the concomitant deletion of the *fis1* and *fis3* genes is not viable in the *L. pneumophila* Philadelphia strain (68). In addition, the roles of Fis1, Fis2 and Fis3 appear to differ between strains. In the Philadelphia strain, Fis1 and Fis3 have been found to directly regulate effector gene expression, whereas Fis2, which we have shown to play a role in the Paris strain, does not (26).

Among the 329 downregulated genes in the Δ*fis1-2* double mutant, approximately one third encode typical transmissive phase traits such as the flagellar regulon, pili-encoding genes and other Dot/Icm independent virulence factors that are predicted to affect the first step of cell invasion (EnhABC and their homologues, RtxA, LidL), but also several regulators involved in the transition between the replicative and the transmissive phase, such as the transmission trait enhancer LetE (bacham and Swanson, 2004), the sigma factors RpoN and FliA (51, 90–92), the RNA chaperone Hfq (93, 94), the two-component system CpxR/A (63, 65, 66), PilS/R and lpp2523/2524 (51), and several secondary messenger cyclic-diGMP metabolizing enzymes known to coordinate cell differentiation, virulence (70, 71) and multicellular behaviour like biofilms (72). Interestingly, other genes previously defined as “late transmission genes” for their strong stationary phase induction, such as the sigma factor-encoding gene *rpoE*, the pilus operon (*lpp0686-0681*), the *enhA*-like gene (*lpp1341*) or the effector genes *sidA* and *sidE* are also strongly down-regulated in the *Δfis1-2* mutant strain. Strikingly, an AT-rich DNA motif, very similar to the Fis binding site consensus sequence, was detected upstream of their respective start codons (51). Our results suggest that this set of late transmission genes is actually directly regulated by Fis regulators. Overall, the *Legionella* Fis proteins appear to regulate, directly or indirectly, transmissive associated traits and thus should be considered as a novel class of transcriptional regulators that govern the biphasic life cycle of *L. pneumophila*. Transcriptomic analysis of *Δfis1*, *Δfis2*, and *Δfis1-2* mutants also demonstrate that Fis regulators control the expression of several *icm/dot* genes that encode the main virulence factor Dot/Icm T4SS. Consequently, the secretion machinery’s functionality is directly affected. This explains, at least in part, the strongly attenuated virulence phenotype of the *Δfis1-2* mutant strain. Little is known about the regulatory factors that control the expression of Dot/Icm-encoding genes, except for CpxR that binds and activates the expression of four *dot/icm* genes (*icmR*, *icmV*, icmW, and *lvgA*)(63, 64). Interestingly, transcriptomic analysis, confirmed by transcriptional fusions, shows that the *IcmPO* (*dotML)*, *icmDJB (dotPNO), icmV-dotA, icmWX* operons as well as *dotV* gene are downregulated while the *dotKJIHGF* (*IcmNMLKEG*) operon is up-regulated in the Δ*fis1-2* double mutant. Importantly, the promoters of these loci include Fis putative binding sites, which suggests that they would be directly controlled by Fis1 and/or Fis2 proteins.

Fis1 and Fis2 not only control the secretion machinery but also regulate the expression of 61 of the 330 genes that encode the Dot/Icm effectors. Specifically, Fis1 and Fis2 repress certain genes at the start of infection (exponential phase) and activate others at later times (post-exponential and stationary phases), thereby contributing to the regulatory network that controls the fine coordination of the expression of Dot/Icm effectors, which includes i) the CpxR/A two component system (TCS) that directly activates or represses about 30 effectors genes(62–66), ii) PmrA/B TCS which activates 42 effector-encoding genes (59–61), and iii) the regulatory cascade LetA/S-RsmY/Z whose role is to relieve CsrA repression in order to activate the expression of about 40 effector genes during the transmissive phase (47, 53). Whether the control of the effector genes is direct or indirect remains unclear, as Fis1 and Fis2 activate the *cpxR/cpxA* and *pmrA/pmrB* genes, as indicated above. Nevertheless, some of the genes repressed by Fis1 and Fis2 that we have identified in this study are those described by Zusman as direct transcriptional targets of Fis1 and Fis3 in the Philadelphia strain, suggesting that at least some of the effector genes are directly controlled by Fis proteins (26).

Although the Fis1 and Fis2 proteins of *L. pneumophila* have limited amino acid sequence similarities to each other and to the canonical Fis protein of enteric bacteria, their quaternary structure is conserved. Both Fis1 and Fis2 bind to an AT-rich region of DNA upstream of their target genes, but Fis1 has a higher affinity for this motif than Fis2. Consistently, Fis1 plays a predominant role in controlling the replicative/transmissive transition, virulence and gene regulation. Importantly, Fis1, Fis2 and even Fis3 are able to interact with each other, suggesting that they are capable of forming heterodimers that could bind with variable affinity to the AT-rich region they target. It is important to note that the three Fis proteins are not produced at the same time or in the same quantities. While large quantities of Fis1 and Fis3, are present at the start of bacterial growth, Fis2 synthesis peaks when the bacteria enter the PE phase. Variations in the intracellular concentration of each Fis protein probably contribute to the sequential regulation of target genes in order to optimize their expression according to the different stages of the *Legionella* life cycle. Consistently, the synergistic effect of the *fis1-2* double mutation on gene expression argues that Fis1 and Fis2 proteins are not simply interchangeable but rather cooperate to modulate gene expression for the benefit of the bacterium. We therefore hypothesize that the duplication of *fis* genes in *L. pneumophila* is not simply a back-up system to compensate for potential deleterious mutations in one *fis* gene, but rather a means to fine-tune the expression of targeted genes, in particular virulence genes.

## MATERIAL AND METHODS

### Bacteria, cells and culture conditions

The *L. pneumophila* wild-type strain used in this work is the strain Paris (CIP 107629T)(95). All the mutant strains, derived from Paris wild-type, are listed in the supplemental material **Table S4**. The *E. coli* strains used in this study are also listed in **Table S4**. Bacterial media, antibiotics and culture conditions were as previously described (69, 96). Axenic *Acanthamoeba* spp. cells were grown on PYG medium (proteose peptone, yeast extract, glucose) at 30°C and *Dictyostelium discoideum* cells were grown in HL5 medium at 22°C. Human monocyte cell line U937 was maintained at 37°C in 5% CO2 and RPMI 1640 medium supplemented with 10% heat-inactivated fetal bovine serum (FBS) (Hyclone, GE Healthcare). Differentiation of U937 monocytes in macrophages was induced with 100 ng/ml phorbol 12-myristate 13-acetate (PMA, Sigma) for 48h.

### Construction and complementation of *L. pneumophila fis* deletion mutants

All the plasmids and primers used in this work are listed in supplemental **Tables S4 and S5**. *L. pneumophila* Paris mutants defective for *fis* genes were constructed by replacing the chromosomal gene *fis1* (lpp0606), *fis2* (lpp1324) or *fis3* (lpp1707) by an antibiotic resistance cassette using double homologous recombination strategy. In brief, an approximately 1-kb fragment was amplified from upstream and downstream of the gene of interest, respectively, and cloned together into the pBluescript KS-II plasmid vector (Stratgene) to generate the plasmids pLPPA33, pLPPA34 and pLPPA35 (**Table S4**). The primers, listed in **Table S5**, were designed such that the gene to be deleted was replaced with an open reading frame consisting of the first and last 6 codons, linked by the *Sal*I restriction site, allowing for the subsequent introduction of an antibiotic resistance cassette. The kanamycin (Kan^R^) or gentamycin (Gm^R^), cassettes were PCR-amplified and inserted into pLPPA33, pLPPA34 and pLPPA35 digested by SalI and blunted with the Klenow fragment (Biolabs), yielding plasmids pLPPA36, pLPPA37 and pLPPA38 (with Kan^R^) and pLPPA48, pLPPA49 and pLPPA50 (with Gm^R^) (**Table S4**). All the constructions were sequenced to verify for correct sequences. Then, for allelic exchange of *fis1*, *fis2* or *fis3*, the entire inserts (the flanking regions and the kanamycin or gentamycin cassette) were PCR-amplified and the resulting DNA fragments were introduced separately into the *L. pneumophila* Paris strain by natural transformation as described previously (68). Recombination events into the chromosome, leading to the replacement of *fis1* or *fis2* by the kanamycin resistance cassette, were selected by plating the cells onto BCYE plates containing the antibiotic and incubating for 4-days at 37°C. The resulting transformants were replica plated and screened for the expected allelic exchanges by PCR amplification and sequencing (**Table S5**). To construct the Δ*fis1-2* double mutant strain, the fragment containing the two flanking regions of *fis1* with the Gm^R^ cassette was PCR-amplified and used to transform the *fis2*-Kan^R^ mutant strain. Transformants were selected by plating cells onto BCYE plates containing both antibiotics.

For complementation experiments, the coding sequence of *fis1*, fis2 and *fis3* with their cognate ribosome-binding site were amplified from genomic DNA of *L. pneumophila* Paris strain and inserted separately into the pMMB207C. The resulting plasmids pLPPA210, pLPPA211 and pLPPA212 (**Table S4**) thus contain the *fis* genes under the control of the promoter *Ptac* that was induced by adding 0.5 nM of isopropyl-β-D-thiogalactopyranoside (IPTG) in the culture medium. For complementation during intracellular infections, *fis1* and *fis2* genes were cloned either separately or together into the pXDC50 encoding the fluorescent protein mCherry to monitor intracellular replication of bacteria (**Table S4**).

### Construction of *E. coli fis* mutants

All of our mutants were derived from BW25113 *E. coli* strain, a common laboratory strain expressing the bacteriophage lambda Red recombination system to perform gene disruptions with double-stranded PCR products (98). The *E. coli fis* gene was either deleted or replaced precisely at the same locus with *fis1*, *fis2* or *fis3* genes of *L. pneumophila* according to the following two-step protocol. First, a cassette containing the toxin-encoding gene *ccdB* under the *araBAD* promoter and a kanamycin resistance gene (patent FR3050997) was PCR-amplified from strain CR201 (**Table S5**) and introduced into the chromosome of BW25113 at the *fis* locus by the λ Red recombinase-mediated gene replacement, generating strain BW25113*fis::kan-pBADccdB* (**Table S4**). Kanamycin-resistant recombinant clones were selected and verified by sequencing, and one of them used for subsequent genetic recombination steps. The *kan-pBADccdB* cassette was then replaced with PCR product corresponding to either an in-frame deleted sequence of *E. coli fis* gene or the coding sequence from start to stop codons of *L. pneumophila fis1*, *fis2* or *fis3*. Ara^R^ colonies were purified and screened for loss of kanamycin cassette, and the *fis* locus was sequenced.

### Intracellular multiplication in host cells

Intracellular growth was performed in human macrophages U937 or in amoebae *A. castelanii* as previously described (71) with some modifications described below. Cells were seeded in 96-well plates at a cell density of 1.10^5^ cells per well and infected with *L. pneumophila* strains harboring M-cherry-expressing plasmids at a multiplicity of infection (MOI) of 1. Infections were carried out in RPMI CO2-independent medium (Gibco) with 5% FBS for U937 cells or in PY special for *A. castelanii,* both supplemented with chloramphenicol and IPTG. After inoculation, plates were centrifuged 10 min at 1500 rpm. Intracellular multiplication was monitored by measuring M-cherry fluorescence every hour with a Tecan Infinite M200 plate reader.

### Endoplasmic reticulum recruitment to the LCV

*D. discoideum* amoebae constitutively expressing a calnexin-GFP fusion were seeded on sterile glass coverslips in 12-well plates in HL5 medium. After one night of adhesion, HL5 was removed and replaced by MB medium (7.15 g.L^-1^ yeast extract, 14.3 g.L^-1^ peptone, 20 mM MES [pH 6.9]) containing M-cherry-expressing bacteria at an MOI=10. The plates were centrifuged at 1500 rpm for 10 min and incubated at 25°C for 90 min. Monolayers were then washed twice with PBS 1X and cells were fixed with 4% paraformaldehyde for at least 30 min. ER-recruitment was observed with an inverted confocal microscopy (Axiovert 200M, Zeiss) with a 63x phase contrast objective lens.

### Total RNAs isolation and purification

Total RNAs were extracted from *L. pneumophila* Paris wt, *fis1*, *fis2* and *fis1-2* mutant strains before RNA-Seq analysis. All experiment were performed in duplicate except for Paris wt and *fis1-2* double mutant for which two independent experiments and sequencing have been done, one in duplicate and the other in triplicate. Briefly, bacterial cultures were incubated at 37°C with shaking in AYE medium until entry in stationary phase (OD ≈5). Then, 10^9^ cells were harvested, pelleted (1 min, 13,000 rpm at 4°C) and immediately stored at -80°C. Thawed cells were resuspended in 50 µL of RNAsnap buffer (18 mM EDTA, 0.025% SDS, 95% formamide) and lysed by incubating the suspension at 95°C for 7 min (protocol adapted from (99). Total RNAs were then extracted by adding 1 ml of TRIzol reagent (Invitrogen) and purified with the Direct-zol^TM^ RNA miniprep kit (Zymmo-research) according to the manufacturer’s instructions. Purified RNAs were eluted in Rnase-free water and quantified with a NanoDrop-2000 spectrophotometer (Thermo Fisher Scientific).

### RNA-seq and differential gene expression analysis

Quality and integrity of RNAs were further evaluated on a fragment bioanalyser before proceeding to rRNA depletion. The cDNA libraries preparation and sequencing were performed at Genewiz/Azenta Life Science (Leipzig, Germany) using Illumina NovaSeq platform with a paired-end 150bp sequencing strategy. Differential expression analysis between the Δ*fis* mutants and the wt Paris strains was performed using an in-house pipeline using Trimmomatic v0.36, Bowtie v2.3.0, HTseq v0.6.1 and the DESeq2 package (100–103) and the resulting *P*-value were adjusted using the Benjamini-Hochberg method. To identify differentially expressed genes, we selected those with a 0,6≥log2FC ≥-0,6 and a *P*-value ≤0,05.

### Construction of luciferase- or gfp-based transcriptional reporter fusions

To construct luciferase-based transcriptional fusions, the 1650-bp firefly luciferase (luc) gene was PCR-amplified from the pLA01 (104) and cloned into the digested KpnI-BamHI pXDC91 vector (105), thus replacing the gfp-reporter gene by luciferase gene to give the pLLA01, a luc-based reporter plasmid. Then, promoter regions, including up to 400-bp upstream and 100-bp downstream of the translational start codon, of the set of genes or operons *lpp0686-0681 (operon pilE), IcmTS* (*lpp0507-0508*), *dotKJIHGF* (*lpp0513-0518*), *icmV-dotA* (*lpp2741-2740*), *dotV* (*lpp0537*), *ralF* (*lpp1932*), *sidG* (*lpp1309*), *marB* (*lpp1761*) and *ankG* (*lpp0469*) were PCR-amplified using primers with HindIII and KpnI restriction sites (**Table S5**) and cloned into HindIII-KpnI digested pLLA01. The resulting plasmids, listed in **table S4**, were sequenced prior to their introduction by electroporation into *L. pneumophila* strains.

To construct substitution mutations on the putative Fis-binding sites of the regulatory regions of *lpp0686*, *ralF* and *IcmT*, site-directed mutagenesis was performed on plasmids pLPPA228, pLPPA229 and pLPPA230 using QuikChange II site-directed mutagenesis kit (Stratagene) as described previously (71). The primers used for mutagenesis are listed in **table S5**. The resulting plasmids named pLPPA228mut, pLPPA229mut and pLPPA230mut, were restrict digested with KpnI-HindIII to excise the mutated-promoter regions that were individually subcloned into pLLA01 cut with the same enzymes. The resulting plasmids, listed in **table S4**, were sequenced prior to their introduction by electroporation into *L. pneumophila* Paris strain. To measure luciferase activity, freshly grown *L. pneumophila* strains carrying the *luc*-reporter fusions were resuspended in AYE containing chloramphenicol and D-luciferin (final concentration 200 µg.mL^-1^) to an OD600 of 0.02. Suspensions were distributed in a black with clear bottom 96*-*well plate (150 μl*/*well) and plate was incubated at 37°C inside a plate reader (Tecan Infinite M200pro) with intermittent shaking to allow bacterial growth. Expression of luc-based transcriptional fusions was monitored by measuring the absorbance at 600 nm (*A*600) and luminescence every 15 min.

### TEM translocation assay

The principle of this assay was described previously (71). Briefly, *L. pneumophila* strains expressing various TEM-β-lactamase fusions proteins (76) were used to infect U937 differentiated-macrophages at an MOI of 10. After 10 min of centrifugation at 1500 rpm, plates were incubated at 37°C with 5% CO2 for 90 min. Then, 20 µL of β-lactamase substrate CCF4/AM substrate was added onto infected cells as described before and plates incubated for an additional two hours at room temperature. When TEM-1-β-lactamase fused to effector is translocated into host cell cytosol, the substrate CCF4/AM is cleaved, and the fluorescence emitted turns from green fluorescence (emission at 530 nm) to blue fluorescence (emission at 460 nm). The translocation efficiency of a specific TEM-effector fusion is expressed as the ratio of the blue fluorescence on green fluorescence. The cytoplasmic enzyme enoyl-CoA reductase (FabI), which is not translocated by the T4SS, is used as a control to validate the translocation assay (76).

### SDS-Page gel electrophoresis and Western blot analysis

Flagellin detection was carried out by SDS-Page gel electrophoresis and immunoblotting using anti-FlaA antibody. Briefly, *L. pneumophila* strains grown in AYE medium to exponential and early stationary phases were lysed by sonication and cell debris were pellet by centrifugation. The protein amount of the supernatant was quantified by Bradford and 1μg of total protein was used for western-blot analysis as described previously (47). For TEM-effector fusion proteins detection, *L. pneumophila* strains were grown on CYE agar plate and resuspended in distilled water. Equal amounts of washed cells (10^9^ cells) were boiled 10 min in Laemmli buffer and 15 µL of each lysate was loaded onto SDS-polyacrylamide gel and run in 0.5X TBE buffer (50 mM Tris base, 50 mM boric acid and 1mM EDTA [pH=8]). Then proteins were electrophoretically transferred to nitrocellulose membrane and subsequently incubated with anti-TEM rabbit serum as a primary antibody and an anti-rabbit peroxidase conjugate as secondary antibody. Immunoblots were revealed with the kit Pierce SuperSignal (Thermo Fisher Scientific).

### Proteomics

Quantitative Mass Spectrometry analyses of three independent cultures of *L. pneumophila* Paris WT grown in AYE liquid medium at 30°C until reaching OD600nm 1, 2, 3, 4, and 5 were done as previously described (96). Protein expression of Fis1, Fis2, and Fis3 were extracted from the obtained dataset (**Table S3**).

### Purification of native Fis proteins

*Escherichia coli* BL21DE3*Δfis* pLysS cells carrying pLLP21 (*fis1*) or pLPP22 (*fis2*) were grown overnight at 37°C in LB broth supplemented with ampicillin and chloramphenicol to maintain both pLysS and pLLP21 or pLPP22 plasmids. Overnight cultures were diluted in LB with antibiotics and grown until an OD600nm of 0.7. At this OD, 1 mM of IPTG was added to cultures and cells were grown for an additional 2h at 37°C to induce overproduction of Fis proteins. Cells were subsequently collected by centrifugation at 4°C and the pellets were frozen at -80°C. Cells were thawed and lysed in 15 mL of cold buffer containing 50 mM Tris (pH 8.0), 5 mM EDTA, 10% glycerol, 1 M NaCl, 1 mM dithiothreitol (DTT) and proteases inhibitors by using the French press. Cell debris were removed by centrifugation at 30,000g for 40 min and the supernatant (approx. 15 ml) was diluted with 35 ml of buffer A (20 mM Tris, 1 mM EDTA, 10% glycerol, 1 mM DTT) and frozen at -80°C. The following purification steps for Fis proteins were performed as previously described (106). Briefly, it consists in a two-step purification protocol combining HiTrap Heparin affinity column first, followed by a cation exchange chromatography (Sulfopropyl column) for Fis2 protein or an anion exchange column (DEAE) for Fis1. At the end of the purification process the cleanest fractions were pooled and dialyzed into Fis storage buffer containing 10 mM Tris (pH 8.0), 1 mM EDTA, 50% glycerol, 5 mM DTT, and 0.5 M NaCl and was stored at -20°C.

### Electrophoretic mobility shift assay

DNA fragments corresponding to the native or mutated promoter regions of *icmT*, *ralF* and *pilE* were amplified by PCR from plasmids listed in Table S1. DNA fragments (250 ng) were incubated with 0, 25, 50, 100 µM of Fis1 or 0, 50, 100 and 200 µM of Fis2. The binding reaction was performed at 25°C for 30 min in a 20-µL volume reaction containing 4 µL of 5X binding buffer (50 mM Tris-HCl [pH=7.5], 7 mM KCl, 1 mM DTT, 5% glycerol and 10 ng/µL Salmon sperm DNA). Samples were then loaded in a 6% native polyacrylamide gel that had been pre-run for one hour at 4°C in 0.5X TBE. Electrophoresis was performed at 100V and the gel was stained with 1 µg/mL of ethidium bromide in 0.5X TBE.

### Bacterial two-hybrid assay

The bacterial adenylate cyclase two hybrid (BACTH) system (Euromedex, France) (107) was used to study protein–protein interactions between *L. pneumophila* Fis1, Fis2, Fis3 or the Fis protein of *E. coli* (Fis_Ec). Each *fis* gene was PCR-amplified with specific primers (Table S2) and introduced into similarly BamHI-digested pKT25 or pUT18C plasmid to generate in-frame gene fusions with Cya T18- and Cya T25-fragment (**Table S4**). Ligation mixtures were introduced into *E. coli* DH5α and transformants were selected on LB plates supplemented with 100 µg/mL Amp (for pUT18C derivatives) or 30 µg/mL Kan (for pKT25 derivatives). All recombinant plasmids were verified by DNA sequencing. Finally, *E. coli* BTH101 (*cya* mutant, Euromedex) cells were co-transformed with a combination of one pUT18C derivative and one pKT25 derivative and the transformants were selected on LB plates supplemented with both Kan and Amp. To analyze interactions between Cya T18- and Cya T25-fusion proteins, two complementary methods were conducted: (1) a screen on LB-X-Gal plates where clones positive for protein–protein interaction turned blue; and (2) a β-galactosidase assay providing a quantitative determination of the efficiency of the functional complementation between pairs of hybrid proteins. Several negative controls were included in each experiment: (1) pUT18C with pKT25 (empty vectors only), (2) pUT18C empty vector with each pKT25-gene fusion, (3) pUT18C-gene fusions with pKT25 empty vector and (3) pUT18C-*lssA* with each pKT25-gene fusion. A positive control (pUT18C-zip with pKT25-zip, provided with the kit) was also included in each experiment. Triplicates were performed for each pair of plasmids. *E. coli* BTH101 cultures are grown overnight at 30°C with shaking in LB supplemented with both antibiotics and 0,5 mM IPTG. 5 µL of cell suspensions were then spotted on LB agar supplemented with ampicillin (100 µg/ ml), kanamycin (50 µg/ml), X-Gal (40 µg/ml), and IPTG (0.5 mM). Plates were incubated at 30°C for 36–48 h under aerobic conditions before being imaged.

### β-galactosidase assay

For β-galactosidase assay, 20 µL of the overnight *E. coli* BTH101 cultures were incubated with 980 µL of Z-buffer (70mM Na2HPO4.12H20, 30mM NaH2PO4.H20, 10mM KCl, 1mM MgSO4.7H20 supplemented with 100 mM β-mercaptoethanol just before use) at 30°C for 5 min. Then, 50 µL of toluene and 0.001% SDS were added and mixed vigorously before being incubated for a further 10 minutes to allow for the lysis of the cells. After letting cell debris settle down at the bottom of the tube, aliquots of 20 µL were transferred into a 96-well flat-bottom microplate containing 180 µl of pre-warm Z buffer. Then, 10 µL of ONPG 0.4 % were dispensed in each well and the enzymatic reaction was carried out at 30°C for 15 min with measurement at OD420nm every 2 min. The relative β-galactosidase activity of each sample was then calculated using Beer-Lambert’s law as followed Activity (mol/L of β-gal produced) = [Δ(OD420 – OD420 in control tube) / Δ min of incubation) x 5000 x dilution factor x optical path (0.9 cm). The factor 5000 in the above formula is the absorption coefficient of o-nitrophenol, without Na2CO3 addition (L/mol/cm). The time points were chosen to be located in the linear part of the kinetic. Results were then expressed as specific activity (mol/L/mg dry weight bacteria) as followed: AS = Activity / bacterial dry weight ∼0.5µg per OD600 unit) considering that 1ml of culture at OD600 =1 corresponds to 500 μg dry weight bacteria. Statistical tests were performed between the different conditions, including a non-parametric Kruskal-Wallis test with a p-value of 0.003669 excluding the positive control, followed with a Dunn’s post-hoc pairwise test without correction carried out on each pair. Significance is represented by stars, where * indicates a p-value < 0.5, ** = 0.1, *** < 0.1 and N.S indicates a p-value > 0.5.

### Phylogenetic analyses to assess the inter- and intra-*Legionella* diversity of Fis paralogs

All available *L. pneumophila* genomes were download from NCBI assembly database (download the May the 30^th^ of 2024). Assembly quality was assessed using Quastv5.0.2 and assemblies with more than 100 contigs were removed of the dataset (108). Species identification was performed using fastANIv1.33, genomes mis-assigned to *L. pneumophila* were removed of the dataset (109). The remaining 5,216 genomes of the dataset were used to generate a local blast database. For phylogenetic analyses on the genus scale, one representative genome of each *L.* non-*pneumophila* was download from NCBI database. Assembly quality was assessed using Quastv5.0.2. *L. tunisiensis* assembly was remove from the dataset due to bad quality. The remaining 64 genomes of the dataset were used to generate a local blast database. tBlastn analyses were conducted against these databases using the amino acid sequences of the three Fis protein from the reference strain *L. pneumophila* strain Paris (CIP107629T) (110). Phylogenetic trees were constructed based on amino acid sequences using seaview v5.0.2 (111).

## Acknowledgments

We thank Virginie Gueguen-Chagnon, Adeline Page and Frédéric Delorme from the Protein Science Facility at the SFR Biosciences (UAR3444/CNRS, US8/Inserm, ENS de Lyon, UCBL), the first for protein purification and the second and third for the Mass spectrometry analyses. We are grateful to Carmen Buchrieser (Pasteur Institute, Paris) for the gift of anti-FlaA antibody. We thank the Dicty Stock Center for *D. discoideum* strains. And finally, we thank Omran Allatif (BIBS, CIRI) for helping with R scripts and statistics.

## Author contributions

E.K. designed research and analyzed the data; E.K., C.A. and J.B. performed all the experimental research except proteomics experiments that were designed, performed and analyzed by L.A. and K.P; C.G. performed the phylogenetic analyses; C.G. and E.K. performed and analyze transcriptomic data. A.V. provided AlphaFold models and analyzed the data. N.B., CR and W.N. provided materials, protocols and technical assistance for confocal microscopy, construction of *E. coli* mutant strains and protein purification, respectively. A.C. contributed to discussions on signaling and virulence control in *L. pneumophila*. E.K., A.V. and P.D. wrote the paper with major input from L.A. and W.N.

